# Chemical signatures delineate heterogeneous amyloid plaque populations across the Alzheimer’s disease spectrum

**DOI:** 10.1101/2024.06.03.596890

**Authors:** Srinivas Koutarapu, Junyue Ge, Maciej Dulewicz, Meera Srikrishna, Alicja Szadziewska, Jack Wood, Kaj Blennow, Henrik Zetterberg, Wojciech Michno, Natalie S Ryan, Tammaryn Lashley, Jeffrey Savas, Michael Schöll, Jörg Hanrieder

## Abstract

Amyloid plaque deposition is recognized as the primary pathological hallmark of Alzheimer’s disease(AD) that precedes other pathological events and cognitive symptoms. Plaque pathology represents itself with an immense polymorphic variety comprising plaques with different stages of amyloid fibrillization ranging from diffuse to fibrillar, mature plaques. The association of polymorphic Aβ plaque pathology with AD pathogenesis, clinical symptoms and disease progression remains unclear. Advanced chemical imaging tools, such as functional amyloid microscopy combined with MALDI mass spectrometry imaging (MSI), are now enhanced by deep learning algorithms. This integration allows for precise delineation of polymorphic plaque structures and detailed identification of their associated Aβ compositions. We here set out to make use of these tools to interrogate heterogenic plaque types and their associated biochemical architecture. Our findings reveal distinct Aβ signatures that differentiate diffuse plaques from fibrilized ones, with the latter showing substantially higher levels of Aβx-40. Notably, within the fibrilized category, we identified a distinct subtype known as coarse-grain plaques. Both in sAD and fAD brain tissue, coarse grain plaques contained more Aβx-40 and less Aβx-42 compared with cored plaques. The coarse grain plaques in both sAD and fAD also showed higher levels of neuritic content including paired helical filaments (PHF-1)/phosphorylated phospho Tau-immunopositive neurites. Finally, the Aβ peptide content in coarse grain plaques resembled that of vascular Aβ deposits (CAA) though with relatively higher levels of Aβ1-42 and pyroglutamated Aβx-40 and Aβx-42 species in coarse grain plaques. This is the first of its kind study on spatial *in situ* biochemical characterization of different plaque morphotypes demonstrating the potential of the correlative imaging techniques used that further increase the understanding of heterogeneous AD pathology. Linking the biochemical characteristics of amyloid plaque polymorphisms with various AD etiologies and toxicity mechanisms is crucial. Understanding the connection between plaque structure and disease pathogenesis can enhance our insights. This knowledge is particularly valuable for developing and advancing novel, amyloid-targeting therapeutics.

## INTRODUCTION

Alzheimer’s Disease (AD) is an age associated disorder affecting 12% over the age of 65 and poses an ever increasing societal and socioeconomic challenge (Matthews *et al*. 2019). The major pathological hallmarks of AD include the progressive, abnormal accumulation and deposition of beta-amyloid (Aβ) peptides as extracellular plaques and hyperphosphorylated tau protein as neurofibrillary tangles (Braak & Braak 1991). The most widely accepted hypothesis suggests that Aβ aggregation and plaque formation are critical early events in AD pathogenesis that initiate a damaging cascade (Hardy & Selkoe 2002) of downstream events such as synaptic changes, tau pathology and eventual neurodegeneration (Selkoe 2002). The relevance of Aβ, and particularly its aggregates in AD, has seen a recent resurgence following positive results from the Phase III study and FDA approval of the Aβ-targeting antibody lecanemab (Hayato *et al*. 2022; Rafii *et al*. 2022; Satlin *et al*. 2016; Swanson *et al*. 2021; EISAI 2022; van Dyck *et al*. 2023) as well as positive Phase III results for donanemab (Sims *et al*. 2023; Mintun *et al*. 2021), which both provide significant support of the amyloid cascade hypothesis (Hardy 1992; Hardy & Selkoe 2002). While amyloid targeting therapies appear to slow down the disease.

However, clinical trial data suggest that patients are different in terms of how well they respond. In detail, all treated patients respond with amyloid red and plaque polymorouction though not all respond well cognitively or in regards to imaging- and fluid-based biomarkers for pathophysiological pathways downstream of Aβ. This suggests that heterogeneity of Aβ plays a role in amyloid pathogenicity.

Amyloid β pathology presents itself through a variety of structural and chemical heterogeneous aggregates including senile plaques as well as cerebral amyloid angiopathy (CAA). Amyloid plaques are observed in the parenchyma and can be categorized as non-fibrillar, diffuse plaques or fibrillar plaques, which includes compact (core only) and classic, cored plaques. The presence of degenerated neurites and axons in the immediate microenvironment of fibrillar plaques denotes those deposits as neuritic plaques. The prevalence of polymorphic Aβ plaque phenotypes is associated with clinical dementia, specifically with fibrillar plaques, where the neocortical load of neuritic deposits correlates with clinical symptoms. In contrast, diffuse plaques are also observed in cognitively unaffected, amyloid-positive individuals suggesting that formation or maturation into fibrillar plaque types is critical in dementia pathogenesis. Consequently, a detailed understanding of the biochemical makeup of distinct amyloid plaque morphotypes is critical to understand the etiology of plaque-driven pathology which is also important in light of amyloid targeting therapies. More so, characterizing vascular Aβ is also important given ARIA occurring in AD immunotherapy, which may be (partly) related to CAA. Commonly, plaque characterization has been based on histological and immune-based tissue staining techniques such as silver staining, Congo red, thioflavin complemented with immunohistochemistry and immunofluorescent microscopy, respectively. All these tools have significant shortcomings with respect to specificity, sensitivity, bias, and throughput that provide solely discrete readout variables that comprehensively form the basis of pathological staging (Montine *et al*. 2012; Mirra *et al*. 1991). This poses significant challenges to understand comprehensively and to quantitatively delineate the biochemical composition of structurally distinct plaques. The advent of novel chemical imaging modalities such as mass spectrometry imaging (MSI) significantly overcome the limitations of biological staining techniques. This approach allows characterization of comprehensive lipid and peptide patterns in situ at 5-10µm resolution (Michno *et al*. 2019c). In the context of AD, our lab has pioneered the application of MALDI MSI for mapping Aβ patterns at the single plaque level in both mouse models (Carlred *et al*. 2016; Michno *et al*. 2021; Michno *et al*. 2020) and postmortem human brain tissue (Michno *et al*. 2019a). Moreover, morphological heterogeneous amyloid pathology can be interrogated using structure sensitive fluorescent probes, such as luminescent oligothiophenes (LCO), that provide both a qualitative and quantitative readout on plaque morphology and amyloid cross β sheet content (Nystrom *et al*. 2013; Rasmussen *et al*. 2017). Herein, we aim to interrogate heterogeneous amyloid plaque pathology with focus on fibrillar plaque morphotypes across different forms of AD. Here we identify specific Aβ patterns of fibrillar and non-fibrillar plaques across different etiological subtypes of AD. We further identified and characterized a distinct subtype of fibrillar plaques, previously described as coarse grain (CG) plaques that given its neuritic nature represents a potential, key phenotype of amyloid toxicity.

## MATERIALS AND METHODS

### Chemicals and Reagents

All chemicals for matrix and solvent preparation were pro-analysis grade and obtained from Sigma-Aldrich/Merck (St. Louis, MO) unless otherwise specified. TissueTek optimal cutting temperature (OCT) compound was purchased from Sakura Finetek (Cat.#: 4583, AJ Alphen aan den Rijn, The Netherlands). Deionized water was obtained by a Milli-Q purification system (Millipore Corporation, Merck, Darmstadt, Germany).

### Patient Samples

Frozen brain tissue samples were obtained from temporal cortex of individuals who had been clinically and pathologically diagnosed with sporadic AD (sAD, n=9) and autosomal dominantly inherited familial AD (fAD, n=6) (Table 1). All cases were obtained through the brain donation program of the Queen Square Brain Bank for Neurological Disorders (QSBB), Department of Clinical and Movement Neurosciences, UCL Queen Square Institute of Neurology. The standard diagnostic criteria were used for the neuropathological diagnosis of AD (Montine *et al*. 2012; Thal *et al*. 2015; Thal *et al*. 2002; Braak & Braak 1991). Ethical approval for the study was obtained from the Local Research Ethics Committee of the National Hospital for Neurology and Neurosurgery, as well as the Ethics Review Board at the University of Gothenburg (Gothenburg, 04/16/2015; DNr 012-15). All studies abide by the principles of the Declaration of Helsinki.

**Table1:** Patient Demographics

| ID | AAO | AAD | Sex | Clinical Diagnosis | Path Diagnosis | Braak | Thal | CERAD | ABC | Mutation |
| --- | --- | --- | --- | --- | --- | --- | --- | --- | --- | --- |
| sAD1 | 43 | 58 | F | AD | AD | VI | 5 | 2 | A3B3C2 |  |
| sAD2 | 54 | 65 | M | AD | AD | VI | 5 | 3 | A3B3C3 |  |
| sAD3 | 58 | 68 | M | PCA | AD | VI | 5 | 3 | A3B3C3 |  |
| sAD4 | 55 | 67 | F | PPA | AD | VI | 5 | 3 | A3B3C3 |  |
| sAD5 | 63 | 79 | M | AD | AD | VI | 5 | 3 | A3B3C3 |  |
| sAD6 | 49 | 69 | F | AD | AD | VI | 5 | 2 | A3B3C2 |  |
| sAD7 | 48 | 63 | M | AD | AD | VI | 5 | 3 | A3B3C3 |  |
| sAD8 | 51 | 62 | F | AD | AD | VI | 5 | 3 | A3B3C3 |  |
| sAD9 | 50 | 66 | F | PPA | AD | vi | 5 | 3 | A3B3C4 |  |
| fAD1 | 42 | 47 | M | FAD | FAD | V | 5 | 3 | A3B3C3 | *PSEN1 A434T & T291A* |
| fAD2 | 46 | 65 | F | FAD | FAD | VI | 5 | 3 | A3B3C3 | *PSEN1 after 200 (R278I)* |
| fAD3 | 35 | 51,9 | F | FAD | FAD | VI | 5 | 3 | A3B3C3 | *PSEN1 Intron 4* |
| fAD4 | 33 | 37 | F | FAD | FAD | VI | 5 | 3 | A3B3C3 | *PSEN1 E120K* |
| fAD5 | 47 | 58 | F | FAD | FAD | VI | 5 | 3 | A3B3C3 | *PSEN1 E184D* |
| fAD6 | 42 | 51 | M | FAD | FAD | VI | 5 | 3 | A3B3C3 | *PSEN1 mutation (Intron 4)* |

AAO: age at onset; AAD: age at death; PCA: posterior cortical atrophy; PPA: primary progressive aphasia; *PSEN1*: Presenilin 1; CERAD: Consortium to Establish a Registry for Alzheimer's Disease; ABC, Combined score based on Thal phase, Braak stage and CERAD Score

### Sample Preparation

For amyloid peptide imaging, we employed a previously validated protocol for robust peptide and protein mass spectrometry imaging (Michno *et al*. 2019). Frozen tissue sections were thawed and dried under vacuum for 15 min. A series of sequential washes of 100% EtOH (60 s), 70% EtOH (30 s), Carnoy’s fluid (6:3:1 EtOH/CHCl_3_/acetic acid) (90 s), 100% EtOH (15 s), H_2_O with 0.2% TFA (60 s), and 100% EtOH (15 s) was carried out.

The sections were then incubated with two LCO fluorophores (tetrameric formyl thiophene acetic acid, q-FTAA, 2.4 µM in MilliQ water and heptameric formyl thiophene acetic acid, h-FTAA, 0.77 µM in MilliQ water), in the dark for 25 min. Subsequently, the sections were subjected to a single 10 min 1x PBS wash and stored in the dark at RT until fluorescent microscopy analysis.

### LCO Microscopy

Multichannel imaging of immuno-stained human brain sections was performed using an automatic widefield microscope (Axio Observer Z1, Zeiss, Germany). Large multi-channel tile scans were captured using EGFP filter sets. All the images were captured using a Plan-Apochromat 20×/0.8 DIC air objective lens.

### Deep learning model for plaque identification

#### Data Collection and Preprocessing

A dataset of LCO based fluorescent amyloid plaque images was collected, ensuring a diverse representation of plaque types. Annotations of the amyloid plaque images were classified into coarse, cored, and diffused classes based on the morphology. The model was trained and tested using a dataset of annotated amyloid plaque images (n = 385) consisting of coarse grain plaques (n = 42); cored plaques (n = 208); diffuse plaques (n = 135). The images were preprocessed to standardize size (120×120), converted to grayscale, and adjusted for contrast.

#### Network Architecture

We designed a fully connected neural network consisting of multiple convolutional layers followed by max-pooling layers for feature extraction. Batch normalization and dropout layers were employed to enhance model generalization and prevent overfitting. The final classification layer utilized softmax activation to output probabilities for each plaque type.

#### Training Procedure

##### Three-fold cross-training

From the 385 images, 20% (n = 78) was held-out as an unseen test dataset. The remaining 307 images were used for training and validation. In each iteration of the cross-training process, two subsets (80% of the data) are utilized for training, while the remaining subset (20% of the data) serves as the validation set. The parameters or predictions of the three trained models are averaged to create a final model. This averaging process helps diminish the influence of specific training-validation splits and enhances the robustness of the final model.

##### Model training

The classification model was trained on each combination of training and validation subsets. This process was repeated three times, covering all possible combinations of training and validation sets. We employed an adam optimizer with a learning rate schedule to efficiently converge towards optimal weights. Data augmentation techniques such as random rotation, flipping, and scaling were applied to augment the training set, improving model robustness and reduce overfitting.

#### Evaluation

##### Evaluation Metrics

The performance of the trained model was evaluated using metrics such as accuracy, precision, recall, and F1-score on the held-out unseen test set (n = 78). Confusion matrices were analyzed to assess the model’s ability to distinguish between different plaque types (Fig S8).

### MALDI MS imaging

Following microscopy, tissues were subjected to formic acid vapor for 20 min. A mixture of 2,5-dihydroxyacetophenone (2,5-DHAP) and 2, 3, 4, 5, 6-pentafluoroacetophenone (PFAP) was used as matrix compound and applied using HTX TM-Sprayer (HTX Technologies LLC, Carrboro, NC, USA). A matrix solution of 5.7 µl/ml of PFAP and 9.1 mg/ml of DHAP in in 70% ACN (Timmers *et al*.), 2% acetic acid / 2% TFA was sprayed onto the tissue sections using the following instrumental parameters: nitrogen flow (10 PSI), spray temperature (75°C), nozzle height (40 mm), eight passes with offsets and rotations, and spray velocity (1000 mm/min), and isocratic flow of 100 μl/min using 70% ACN as pushing solvent.

MALDI-MSI experiments were performed on a rapifleX Tissuetyper MALDI-TOF/TOF instrument (Bruker Daltonics) using the FlexImaging and FlexControl (v5.0, Bruker Daltonics) software. Measurements were performed at 10μm spatial resolution, at a laser pulse frequency of 10 kHz with 200 shots collected per pixel. Data were acquired in linear positive mode over a mass range of 1500–6000 Da (mass resolution: m/Δm=1000 (FWHM) at m/z 4512). Pre-acquisition calibration of the system was performed using a combination of peptide calibration standard II and protein calibration standard I (Bruker Daltonics) to ensure calibration over the entire range of potential Aβ species.

### MSI Data Analysis

Following MALDI imaging spatial segmentation was used by HCA/ bisecting k mean clustering to identify the localization pattern associated with single plaques. The HCA-derived pseudo clusters revealed distinct plaque signatures. LCO stained plaques were identified from the tile scan images obtained from fluorescent widefield microscopy prior to MSI analysis. These plaques were identified across the whole cortical brain sections (approximately 1 cm^2^) from each patient involved in the study. These tile scans were aligned onto the single ion images obtained from MSI. Segmented MSI cluster images were co-registered with LCO stained images to identify plaque polymorphs as identified by DL.

For statistical analysis, regions comprising of annotated plaques were marked as ROIs in FlexImaging (v5.0, Bruker Daltonics). Data from 5-10 plaques per subtype across all patient samples within each group were exported as *.CSV files and imported into Origin (version 8.1 OriginLab, Northampton, MA). Bin borders were generated using the PeakAnalyser function in Origin. Peak bin were values were obtained using an in-house developed R script as described previously (Hanrieder *et al*. 2011). All the data were compiled in MS Excel.

Average peak data from the different plaque-ROI were analyzed by grouped, univariate statistical comparisons of the respective plaque subtypes across patients by means of paired t-statistics (p<0.05) and also Holm-Šídák’s post hoc multiple comparisons test using Prism (v.9, GraphPad, San Diego, CA, USA). Metaboanalyst was used to perform multivariate analysis.

### Immunohistochemistry

For immunohistochemistry and fluorescent amyloid imaging, sections on super frost glasses were fixed in gradient concentration of ice cold 95%, 70% ethanol and 1XPBS at room temperature. Sections were then blocked with (Bovine Serum Albumin BSA, Normal Goat Serum NGS, Triton in 0.1% PBST) for a period of 90 minutes at room temperature. The sections were then incubated with cocktail of two primary antibodies PHF-1 (Courtesy of Dr. Peter Davies, Feinstein Institute for Medical Research) and RTN3 (Catalog # ABN1723) or 6E10 (anti-amyloidβ 1-16) (Catalog # 803003) (diluted in PBS-T with 0.2% NGS (6E10: 1:500, PHF-1: 1:500 and RTN3: 1:1000) for over 18 hours at 4°C. Sections were washed with 0.1% PBST and incubated with secondary antibodies (Alexafluor594 Catalog # A32740 and/or Alexafluor647-Catalog # A32728) for a duration of 60 min at room temperature. All the brain sections were then treated with autofluorescence quenching agent TrueBlack™ 1X for 30 sec and were later subjected to three 1XPBS washes of 5 minutes each. To stain amyloid and tau morphologies, antibody-stained sections were incubated with two previously validated LCO fluorophores (tetrameric formyl thiophene acetic acid, q-FTAA, 2.4 µM in MilliQ water) and heptameric formyl thiophene acetic acid (h-FTAA, 0.77 µM in MilliQ water), in the dark for 25 min. Subsequently, the sections were subjected to a single 10 min 1x PBS wash followed by mounting with DAKO fluorescent mounting media and incubated in the dark for 24 hours until further imaging.

The multichannel imaging of immuno-stained human brain sections was performed using an automatic widefield microscope (Axio Observer Z1, Zeiss, Germany). Large multi-channel tile scans were captured using EGFP, Alexafluor594 and Alexafluor647 filter sets. All the images were captured using Plan-Apochromat 20×/0.8 DIC air objective lens. The acquisition settings were adjusted to prevent saturation or bleed through during the acquisition of multichannel images acquired with EGFP/Alexafluor594 / Alexafluor647 filter sets.

FIJI ImageJ was used for post processing of images. Here, files from each channel were split into gray scale images and were subjected to background subtraction. Here EGFP/Green channel with LCO stained plaques are used as a reference image for marking the regions occupied by plaques. Segmentation was performed on the green channel images using Li threshold method. Wand tool was used to mark the segmentation and was saved to ROI manager. These ROI annotations were used to acquire intensity measurement from the corresponding gray scale image of anti PHF-1(Alexafluor594) channel and anti RTN3 (Alexafluor647) channel.

Fluorescent intensity circumventing the plaques at 5µm interval (upto 35µm) was measured using an inhouse built macro. Here 3 channel image with qhFTAA (EGFP) channel, PHF-1 (Alexafluor594) channel, RTN3 (Alexafluor647) channel was used extract intensity values. Obtained data was compiled in MS excel. Fluorescence intensities were calculated as intensity/µm^2^.

## Results

### 1. Correlative chemical imaging and deep learning identify polymorphic plaque phenotypes across different clinical pathologies

Herein we set out to delineate amyloid peptide signatures of polymorphic plaque type across sporadic and familial forms of AD (sAD and fAD).

For this we implemented a multimodal chemical imaging strategy based on MALDI MSI and fluorescent microscopy using functional amyloid probes (LCO) (Michno *et al*. 2019a). The segmentation analysis revealed characteristic chemically diverse localization patterns (pseudoclusters) that resemble plaque features across the brain tissue sections. Alignment of the HCA segmentation map with the LCO amyloid microscopy allowed for tentative identification of different plaque morphologies (plaque types) that align with distinct pseudoclusters in the HCA. This in turn revealed the corresponding amyloid peptide makeup encoded in the MALDI MSI data space. In sAD, this LCO/MSI approach identified three clusters of plaque types, including diffuse plaques (DP) and cored plaques (CP). This further identified an isolated subcluster of fibrillar plaques (Schmidt *et al*. 1995; Dickson & Vickers 2001), previously referred to as coarse grain phenotype (Boon *et al*. 2020). In fAD tissue, two main clusters were identified comprising only fibrillar plaques i.e. cored plaques and coarse grain plaques. In one patient, prominent cotton wool plaque (CWP) pathology was observed and could be identified as separate plaque cluster in the MALDI data (Fig 1C).

**Figure 1:**
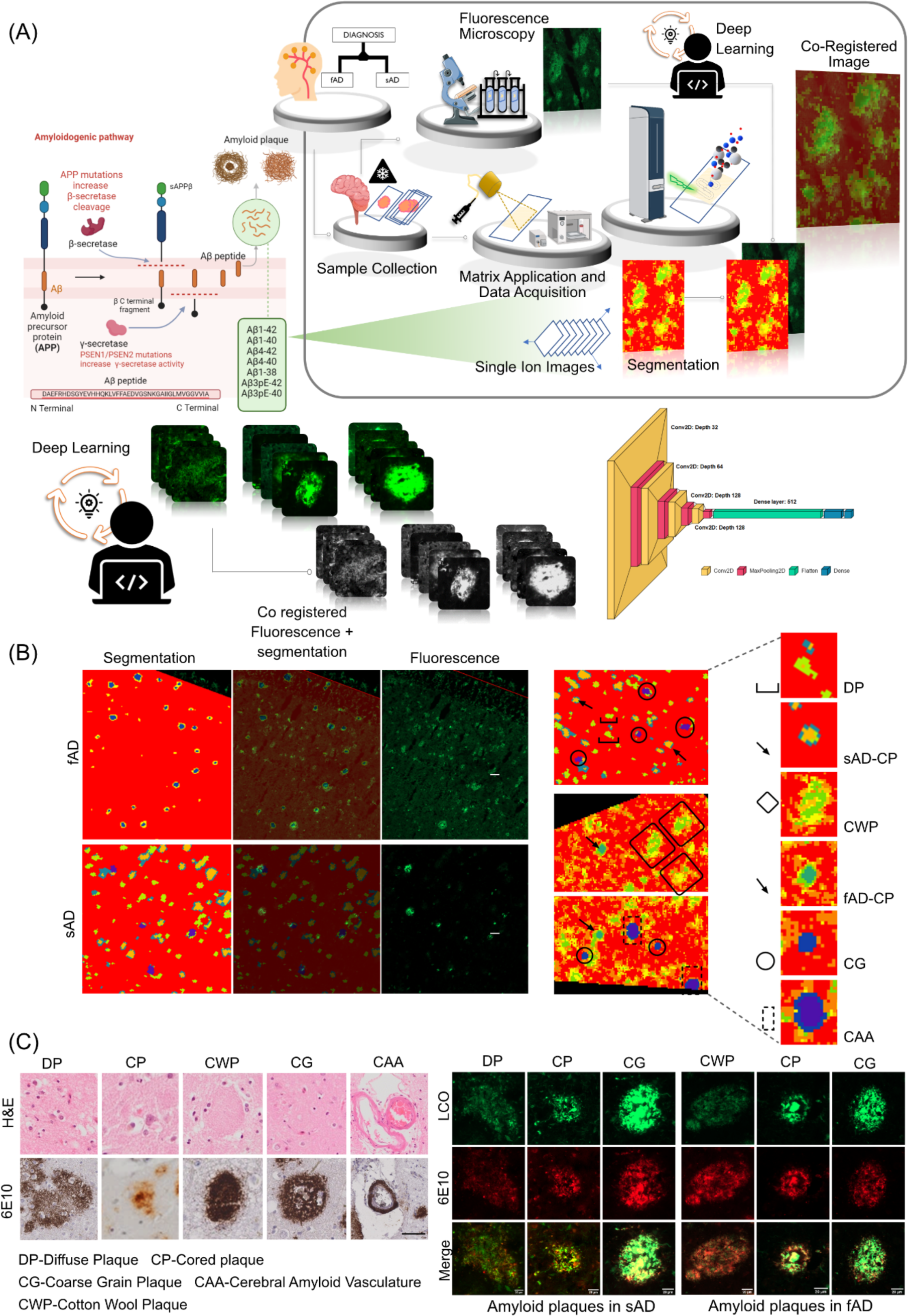
Spatial segmentation of the matrix assisted laser desorption/ionization mass spectrometry (MALDI-MS) imaging data from individuals with sporadic Alzheimer’s disease (sAD) and familial AD patients (fAD). (B) Regions of interest marking qhFTAA based fluorescent amyloid staining of sections from sAD and fAD brain tissues. Registration of fluorescent images with pseudoclusters obtained through spatial segmentation of MALDI MS imaging data. Alignment of amyloid plaques with the pseudoclusters indicates the presence of different plaque morphologies. Spatial segmentation of MALDI MS imaging data and. (C) Visualization of diffuse plaque, cored plaque and coarse grain plaque in sAD and cotton wool plaque, cored plaque and coarse grain plaque in fAD cases through LCO/ antiAβ(6E10) co staining. Scale bar: 20µm. Visualizations of diffuse plaque, cored plaque, cotton wool plaque, coarse grain plaque and amyloid vasculature through H&E and antiAβ(6E10) staining. Scale bar: 40µm

Co-registration of LCO microscopy plaque images onto MALDI MSI data for classification of plaques into different morphologies is subject to bias. To address this issue, we developed a deep learning model to classify these plaques automatically. This model was employed on MALDI-MSI co-registered LCO images, which allowed us to capture plaque specific peptide signatures from MALDI-MSI. In the confusion matrix, with a total of 78 instances, the model correctly classified 72 instances correctly indicating a strong ability to distinguish between the three classes (Fig. 2). However, the model also misclassified 6 instances as suggesting some level of error in the classification into all three classes. Despite these misclassifications, the overall accuracy remains high at 92.3%, reflecting the model’s proficiency in classifying instances correctly. Additionally, the recall of 0.90 (coarse), 0.95 (cored), 0.88 (diffused) suggests the model effectively captures most true positive instances and precision values of 0.82 (coarse), 0.91(cored), and 1 (diffused) indicates a low rate of false positive predictions. The F1 score provides a balance between precision and recall and is useful when the classes are imbalanced, as it is for this dataset. An F1 score (0.86: coarse, 0.93: cored, 0.94: diffused) closer to 1 indicates perfect precision and recall, while a score closer to 0 reflects poor performance. Our CNN-based approach achieved state-of-the-art performance in classifying amyloid plaque images into coarse, cored, and diffused categories. The model demonstrated high accuracy and robustness across diverse plaque morphologies. Furthermore, qualitative analysis revealed the model’s capability to capture intricate features associated with each plaque type.

**Figure 2:**
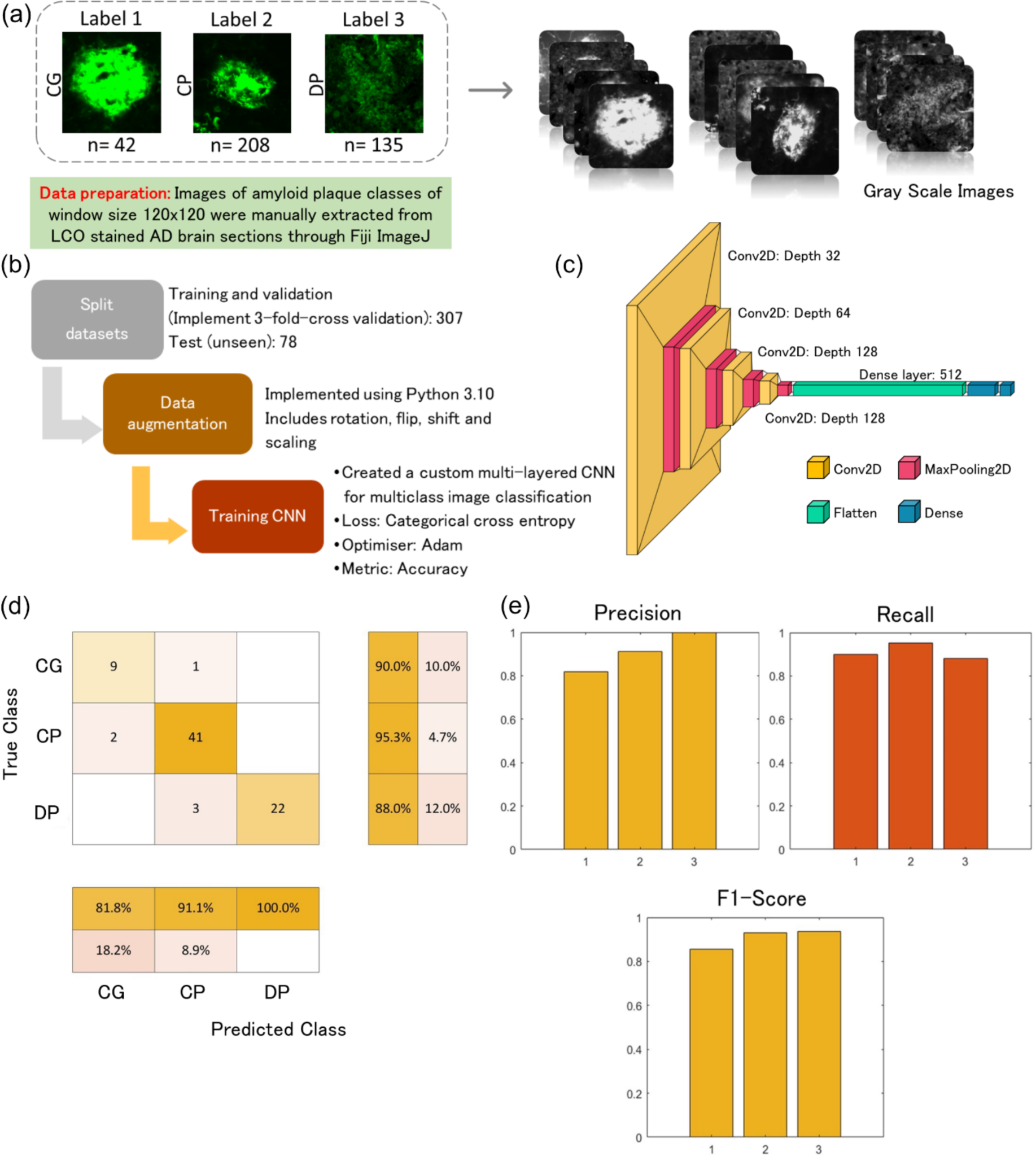
Plaque classification using deep learning. A) Data preparation and preprocessing. Labelled images were manually extracted, categorised, and resized to a standard size (120×120). The images were converted to grayscale and adjusted for contrast. B) Model training and development. The datasets were split into training (n= 307) and test sets (n = 78). Training datasets were split further for training and validation to execute three-fold cross validation and models were trained using various training parameters. The final model was obtaining by averaging the models from three-fold cross validation D) Network architecture E) Results of evaluation using standard classification metrics. The final model achieved an accuracy of 92.3% in the unseen test set (n = 78).

### 2. Amyloid fibrillization is associated with increased Aβ1-40 deposition

Following the DL based classification, the first aim was to study peptide signatures associated with different plaque morphologies in brain tissue across different AD etiologies. In AD brain tissue, senile plaques comprise both diffuse and cored deposits. Further, a variant of diffuse plaque phenotypes called cotton wool plaques, are observed in some fAD cases. Of note, the presence of neuritic, fibrillar plaques has been linked to cognitive symptoms and disease progression (REF). This suggest that plaque maturation from diffuse into cored and eventually neuritic plaques is critically associated with the development cognitive symptoms and AD respectively and that the molecular basis for plaque maturation is reflected in their Aβ signature. To approach this, MS spectral data extracted for diffuse- and cored plaques in sAD, as identified by DL, were compared by means of multivariate analysis (OPLS-DA) and follow up univariate group statistics.

First, OPLS-DA based multivariate analysis of the plaque spectral data separated cored and diffuse plaque types in sAD based on their Aβ signatures (Fig. 3A-aI). The primary loadings that underlie those separations (VIP scores) reveal then the biochemical, i.e., Aβ-, signature associated with the respective plaque types. Here, the OPLS-DA derived VIP scores show Aβ1-40 and Aβ1-42 along with N-terminally truncated, pyroglutamated forms (pE) of Aβx-42, i.e., Aβ3pE-42, Aβ11pE-42 as the critical peptides discriminating diffused plaques from cored plaques in sporadic AD(Fig.3A aII). Similarly, the clustering patterns retrieved from the peptide intensities, as visualized by heatmaps, separate cored and diffuse plaques (Fig.3A bI, bII). Here the main clusters showed distinctly higher intensities for Aβ1-40, Aβ1-42, Aβ11pE-42, Aβ3pE-42 in CP as compared to DP (Fig. 3A c).

**Figure 3:**
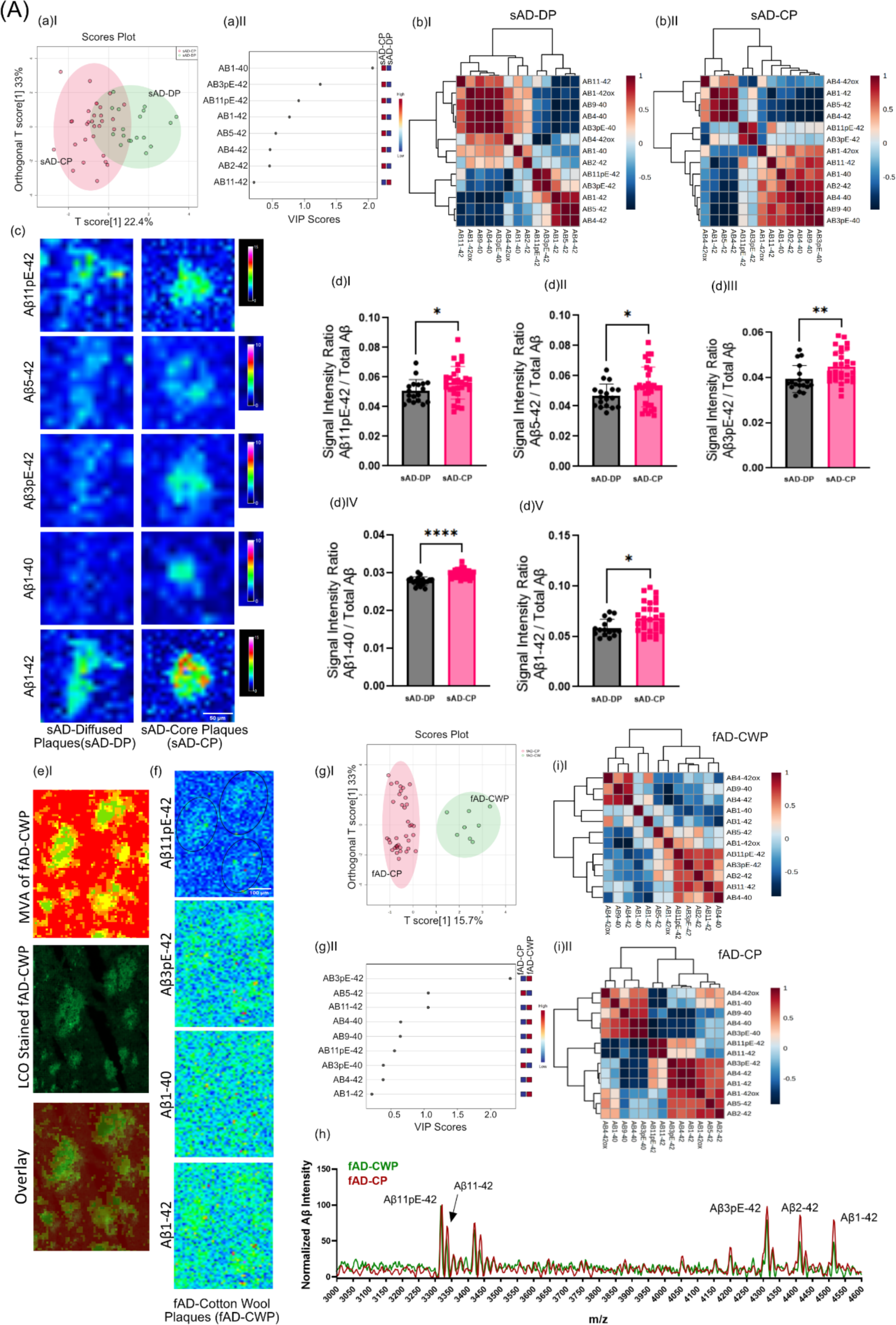
(A) Differences in between fibrillar and non-fibrillar plaques in sAD and fAD. (a I) OPLS-DA separating sAD-DP and sAD-CP (a II) VIP scores indicating peptides which separates sAD-DP from sAD-CP plaques. (b) Clustered intensity heatmap from the individual plaques indicate overlapping separation of sAD-DP and sAD-CP population (OPLS model characteristics: R2X-0.224; R2Y-0.471; Q2-0.44)(c) MALDI MSI single ion images of amyloidβ (Aβ) peptides. Scalebar: 50µm. (d)I-V univariate group statistics for signal intensity of Aβ species normalized to total Aβ (number of patients: n=9 (sAD), number of plaques per type and patient N=4-6). Characterization of Cotton Wool Plaques. (e) HCA based image analysis of fAD brain tissue (red) together with LCO imaging (green) identifies CWP associated MSI profiles and (f) Single ion images. g(I-II) Clustered intensity heatmap from the individual plaques indicate overlapping separation of fAD-CWP and fAD-CP population (OPLS model characteristics: R2X-0.157; R2Y-0.737; Q2-0.604) (h) average mass spectra show relatively higher degrees of pyroglutamated species as compared to Aβ1-42. Pearson’s correlation matrix of Aβ peptides in (i)I fAD-CWP, (i)II fAD-CP Univariate analysis of individual plaque data showed significantly higher levels of Aβ11pE-42(p<0.05), Aβ3pE-42 (p < 0.005), Aβ1-40 (p<0.001) along with Aβ1-42 (p < 0.05) in CP as compared to DP (Fig. 3A dI-V), while other VIP such as Aβ4-42 were not statistically different in between CP and DP.

This suggests that plaque maturation and fibrilization in sAD is predominantly associated with deposition of Aβ1-40 as well as increased deposition of Aβ1-42 and its corresponding pyroglutamated isoforms Aβ3pE-42 and Aβ11pE-42.

We further, expanded this analysis to fAD though only non-fibrillar plaques were observed within one fAD individual (with a double substitution in *PSEN1*; A434T & T291A) showing a morphological phenotype attributed as cotton wool plaques (CWP) that could be delineated on their MSI profiles and validated by correlative LCO microscopy (Fig 3B) (Ryan *et al*. 2016). OPLS-DA based MVA segregated fAD CWP and fAD CP. VIP scores suggest Aβ3pE-42 as the critical peptide discriminating fAD-CWP and fAD-CP (Fig 3B aIII-IV). Also, the clustering patterns retrieved from the peptide intensities, as visualized by heatmaps, separate fAD-CWP and fAD-CP (Fig 3B aV-VI). The MSI patterns of CWP was similar to cored plaques in fAD, showing prominent levels of Aβ11pE-42, Aβ3pE-42, Aβ2-42, Aβ4-42 and Aβ1-42 along with small detectable amounts of Aβ1-40. Here, the relative content of Aβ11pE-42/ Aβ1-42 and Aβ3pE-42/ Aβ1-42 was higher in CWP than in cored plaques (Fig. 3B a VII)).

### 4. Fibrillar plaques in sAD and fAD brain show different content of dystrophic and axonal neurites

Following the initial analysis on cored and diffuse plaques, we then further investigate the fibrillar plaque phenotypes, observed both in sAD and fAD. Interestingly, our HCA based image segmentation of the LCO/MSI data revealed two subtypes of fibrillar plaques annotated as (classic) cored plaques and coarse grain type plaques in line with previous results (Fig. 1) (Boon *et al*. 2020).

Using immunohistochemical and morphological characterization offibrillar plaques in sAD and fAD brain tissue, we investigated the relation of those different plaques with axonal and dystrophic neurites as indications for neurotoxicity. Here we make use of the mirrored section setup, where the facing surface of consecutive sections allowing for correlative MSI and IHC. Plaques including cored- and coarse grain plaques as well as diffuse and CWP were visualized with LCO probes that allow to stain both mature and prefibrillar amyloid aggregates and allow to outline the characteristic plaque morphologies. Immunohistochemistry towards paired helical filament Tau (PHF-1) was used to characterize neuritic content within different plaque morphotypes (Fig. 4A-B, Table 2). Similar to the analysis of diffuse and cored plaques, we first compared neuritic vs non-neuritic, cored plaques in sAD. Here we observed a PHF-1 positive population among cored plaques in addition to coarse grain plaques (Fig. S4a). We therefore first investigated the difference in amyloid peptides among cored and neuritic, cored plaques using OPLS-DA (Fig. S4b). Here, plaques that stain positive for PHF-1 showed significantly higher levels of Aβ1-42ox and Aβ2-42. These findings are well line with previous data obtained from a smaller cohort (Fig. S4 c, d). We then expanded these analyses towards coarse grain plaques in both sAD and fAD brain tissue. Similarly, LCO patterns were used to outline plaque types along with PHF-1 for characterizing axonal neurites within plaques. Further, RTN3 immunolabelling was performed to stain for dystrophic neurites. Here, coarse grain plaques showed prominent staining for PHF-1 as compared to cored plaques (Fig. 4A, B). Similarly, to PHF-1, RTN3 showed increased staining in coarse grain plaques compared to cored plaques (Fig 4. C, D). Interestingly, we did not notice prominent PHF-1/RTN3 positive dystrophy associated with diffuse plaques in sAD and CWP in fAD cases. Together these data indicate that coarse grain plaques are more neuritic in nature than cored plaques.

**Figure 4.**
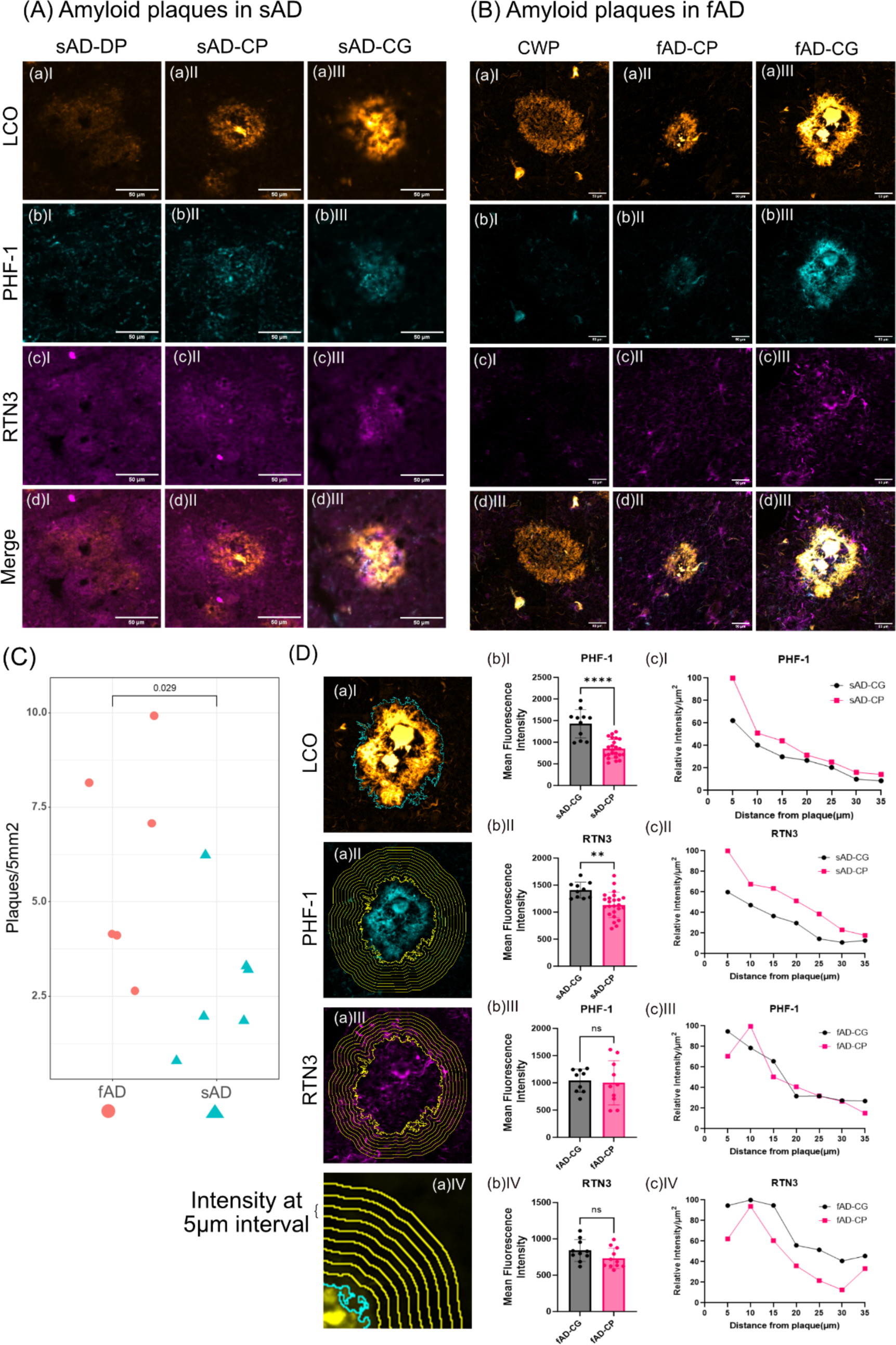
Axonal and dystrophic neurites in Aβ plaque pathology. Combination of two different Luminescent Conjugated Oligothiophenes (LCOs), namely Quadro-formyl thiophene acetic acid (q-FTAA) and Hepta-formyl thiophene acetic acid (h-FTAA) is employed to stain different amyloid polymorphs like (A) diffuse plaques, cored plaques and coarse grain plaques. LCOs stain premature amyloid fibrils in the (a)I diffuse plaques, (a)II densely packed matured amyloid fibrils in the core of the cored plaques and premature fibril in the corona of the cored plaque and (a)III a relatively homogenous distribution of the amyloid fibrils across the coarse grain plaques in sAD. (b) Localization of the PHF-1 staining of the same plaques indicates almost no to very feeble colocalization in diffuse and in cored plaques while a prominent colocalization of qhFTAA with PHF-1 is observed in the case of coarse grain plaques. (c) Similar RTN3 indicates dystrophic neurites. (d)I Mean fluorescent intensity of d(I) RTN3 in sAD-CG and sAD-CP (d)II PHF-1 in sAD-CG and sAD-CP. (B) Similarly, (a)I cotton wool plaques, (a)II densely packed matured amyloid fibrils in the core of the cored plaques and premature fibril in the corona of the cored plaque and (a)III coarse grain plaque. (b) Localization of the PHF-1 staining of the same plaques indicates almost no to very feeble colocalization in cotton wool plaques and in cored plaques while a prominent colocalization of qhFTAA with PHF-1 is observed in the case of coarse grain plaques. (c) Similarly, RTN3 indicates dystrophic neurites. (C) Higher plaque density (No. of plaques/5mm^2^) in fAD than sAD. (D) Fluorescent intensity profile circumventing the plaques was measured from CG and CP from both sAD and fAD cases. These value were captured from (b) PHF-1 (Alexafluor594) channel, (c) RTN3 (Alexafluor647) channel (d) at 5µm interval (up to 35µm). Mean fluorescent intensity of the plaque occupied area for (b)I PHF-1 in sAD-CG and sAD-CP, (b)II RTN3 in sAD-CG and sAD-CP, (b)III PHF-1 in fAD-CG and fAD-CP and (b)IV RTN3 in fAD-CG and fAD-CP. and relative change of intensity at (D aIV) 5µm interval away from the plaque for (c)I PHF-1 in sAD-CG and sAD-CP, (c)II, RTN3 in sAD-CG and sAD-CP, (c)III, PHF-1 in fAD-CG and fAD-CP and (c)IV RTN3 in fAD-CG and fAD-CP. (Univariate analysis: *p<0.05, **p<0.01, ***p<0.001, ****p<0.0001).

**Table 2:**
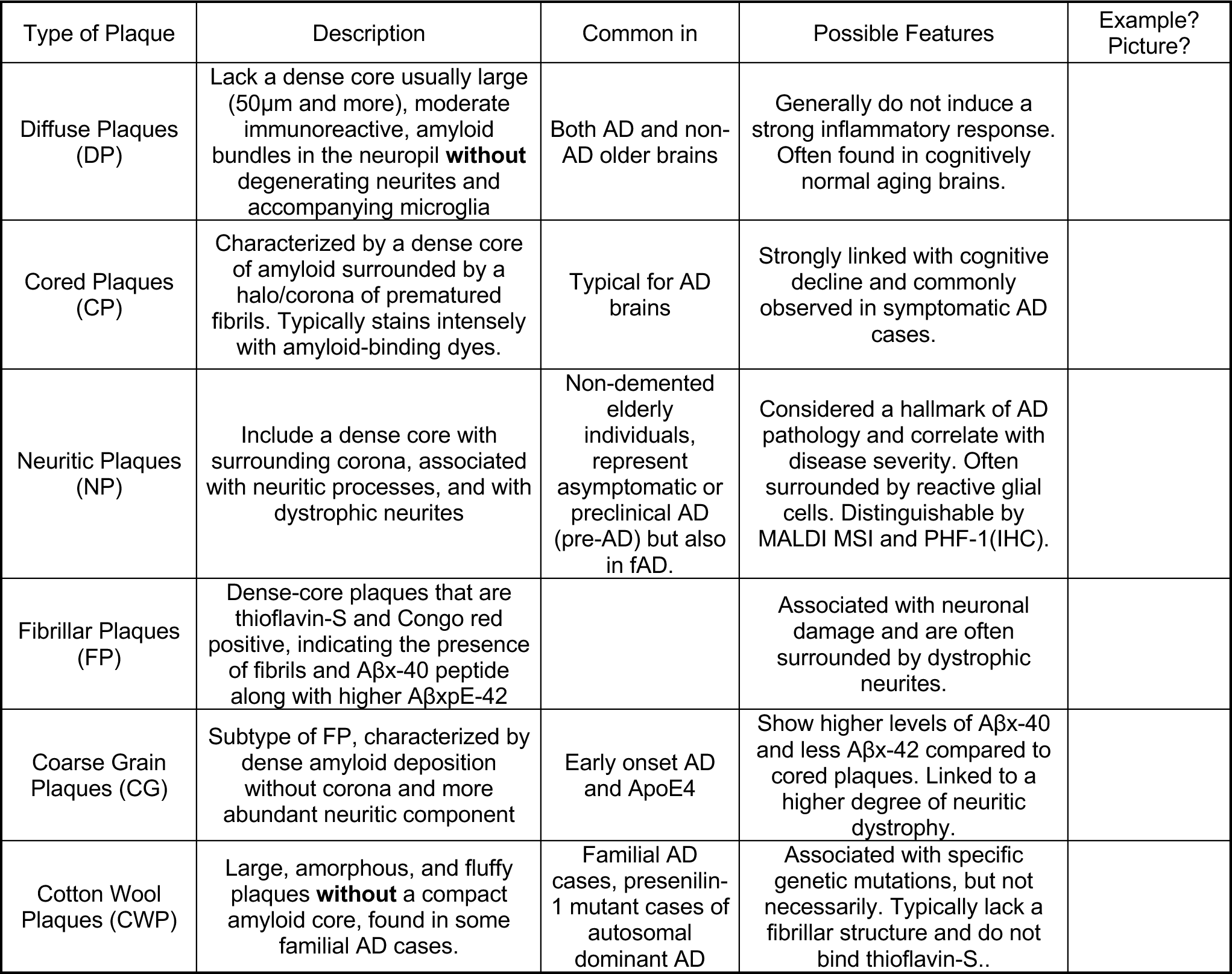
Summary of features associated with various plaque morphotypes

| Type of Plaque | Description | Common in | Possible Features | Example?<br>Picture? |
| --- | --- | --- | --- | --- |
| Diffuse Plaques (DP) | Lack a dense core usually large (50µm and more), moderate immunoreactive, amyloid bundles in the neuropil **without** degenerating neurites and accompanying microglia | Both AD and non-AD older brains | Generally do not induce a strong inflammatory response. Often found in cognitively normal aging brains. |  |
| Cored Plaques (CP) | Characterized by a dense core of amyloid surrounded by a halo/corona of prematured fibrils. Typically stains intensely with amyloid-binding dyes. | Typical for AD brains | Strongly linked with cognitive decline and commonly observed in symptomatic AD cases. |  |
| Neuritic Plaques (NP) | Include a dense core with surrounding corona, associated with neuritic processes, and with dystrophic neurites | Non-demented elderly individuals, represent asymptomatic or preclinical AD (pre-AD) but also in fAD. | Considered a hallmark of AD pathology and correlate with disease severity. Often surrounded by reactive glial cells. Distinguishable by MALDI MSI and PHF-1(IHC). |  |
| Fibrillar Plaques (FP) | Dense-core plaques that are thioflavin-S and Congo red positive, indicating the presence of fibrils and Aβx-40 peptide along with higher AβxpE-42 |  | Associated with neuronal damage and are often surrounded by dystrophic neurites. |  |
| Coarse Grain Plaques (CG) | Subtype of FP, characterized by dense amyloid deposition without corona and more abundant neuritic component | Early onset AD and ApoE4 | Show higher levels of Aβx-40 and less Aβx-42 compared to cored plaques. Linked to a higher degree of neuritic dystrophy. |  |
| Cotton Wool Plaques (CWP) | Large, amorphous, and fluffy plaques **without** a compact amyloid core, found in some familial AD cases. | Familial AD cases, presenilin-1 mutant cases of autosomal dominant AD | Associated with specific genetic mutations, but not necessarily. Typically lack a fibrillar structure and do not bind thioflavin-S.. |  |

The IHC experiments were further expanded towards fAD brain tissue. Here, most prominently cored plaque and coarse grain plaque pathology was observed in all patient samples along with CWP for one patient (Fig.4B). No PHF-1- or RTN3-positive neurites were observed in CWP. In contrast both CP and CG plaques in fAD showed positive staining for PHF-1 and RTN3 i.e. neurites and dystrophic neurites. Here, similarly to sAD, coarse grain plaques showed more pronounced neuritic staining as compared cored plaques (Fig 4B).

Together this suggests that coarse grain plaques are a relevant neuritic plaque population. The absence of non-neuritic cored plaques in fAD further supports that pathology onset relates to the presence of fibrillar plaques with neuritic profiles including both PHF-1/Tau positive neurites and dystrophic neurites highlighted by RTN3. In line with this we observe a higher plaque load of CG in fAD that also show earlier onset of disease (Fig.4C).

Our analysis to understand the difference between sAD and fAD plaques revealed significantly higher RTN3 and PHF-1 intensity values for sAD plaques than with fAD plaques. We measured the mean fluorescence intensity of RTN3 and PHF-1 beneath LCO stained plaque area.

Results of which indicate that the fluorescence intensity of (Fig. 4wD bI) PHF-1 and (Fig. 4D bII) RTN3 is higher in sAD-CG than sAD-CP, while no such difference is seen between fAD-CG and fAD-CP (Fig. 4D bIII, IV). We further measured RTN3 and PHF-1 intensity profile circumventing the plaques at consequent 5µm wide concentric circles up to 35µm away from the plaque (Fig.4D bII, IV) These results suggest that the relative intensity profile of RTN3 and PHF-1 around the sAD-CP is relatively higher than that of sAD-CG. In the case of fAD-CG and fAD-CP intensity of RTN3 was relatively higher than fAD-CP and the PHF-1 intensity profile decrease accordingly (Fig.4D bVI, VII). For all the cases the relative intensity profile of RTN3 and PHF-1 decreased as the distance from the plaque increased.

### 5. Coarse grain plaques are characterized by high Aβ x-40 deposition

Following the immunohistochemical and morphological characterization of fibrillar plaques we then interrogated the Aβ profiles associated with these plaque populations as revealed through MALDI MSI. The comprehensive Aβ patterns allowed us to identify the plaque types through hierarchical cluster analysis (HCA)-based image analysis (Fig. 1).

Moreover, comparative group comparison through multivariate OPLS-DA did reveal a small separation in between cored plaques in fAD and sAD (Fig. 5A, B). HCA of the peptide data intensity patterns showed separation into two main clusters.

**Figure 5.**
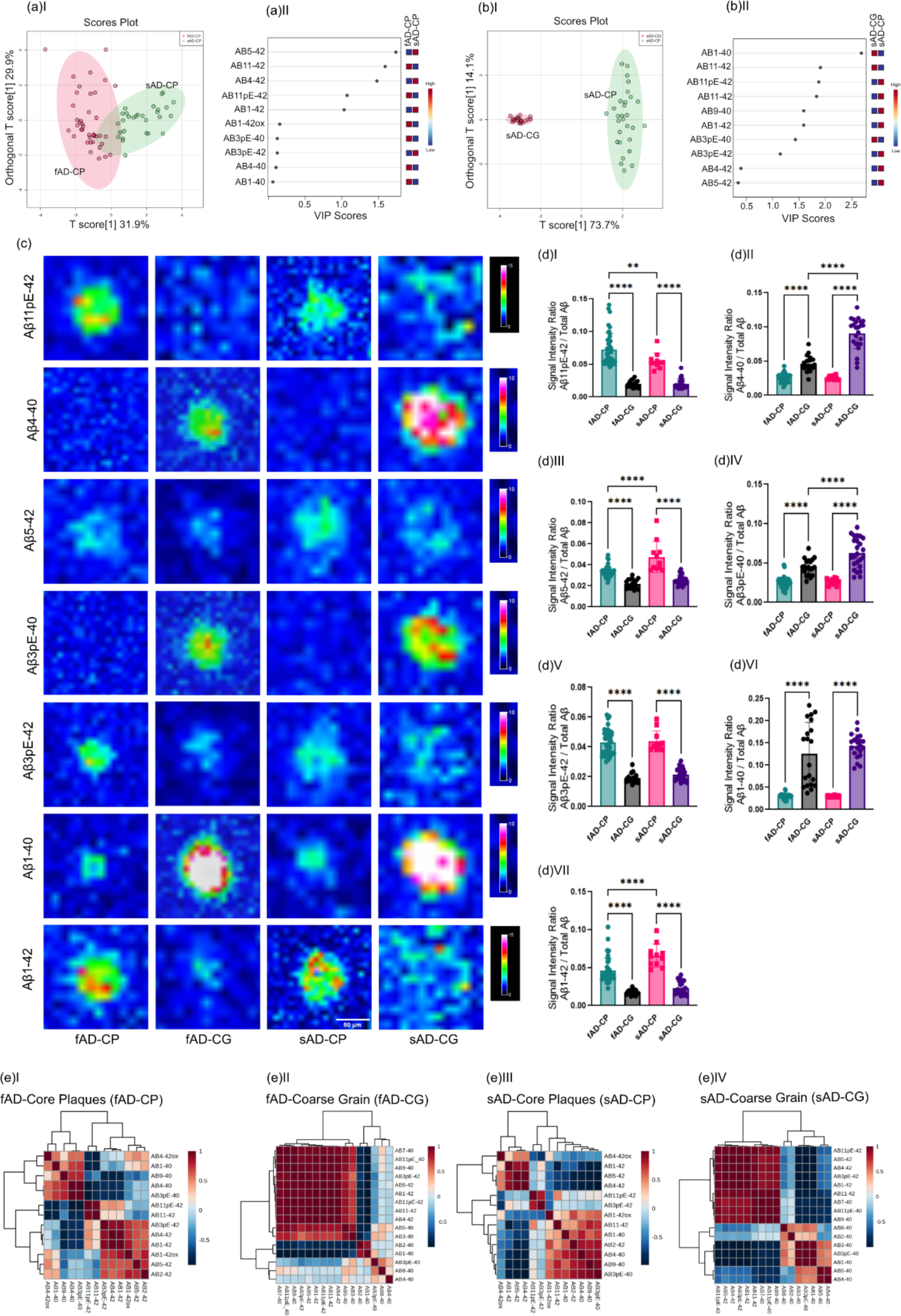
Comparison of peptide patterns in fibrillar plaques: (aI) OPLS-DA of fAD-CP sAD-CP plaques. indicates the partial separation of fAD CP and sAD CP with (OPLS model characteristics: R2X-0.319; R2Y-0.624; Q2-0.609). (aII) VIP scores. (b I) OPLS-DA of sAD-CP and sAD-CG plaques show strong separation (OPLS model characteristics: R2X-0.737; R2Y-0.948; Q2-0.947). (bII) VIP scores. (c) Single ion images of amyloidβ (Aβ) peptides. (d) Corresponding box plots showing univariate comparison of single ion intensities with respective plaque types. (e) Pearson’s correlation matrix of Aβ peptides in (e)I fAD-CP, (e)II fAD-CG, (e)III sAD-CP (e)IV sAD-CG plaques.

Together, OPLS-derived VIP and univariate analysis showed higher levels of Aβ11pE-42 (p < 0.05) in fAD cored plaques, while Aβ5-42 (p < 0.05), and Aβ1-42(p < 0.0001) were found to be higher in sAD cored plaques (Fig. 5C, D).

No significant difference was observed for Aβ3pE-42, Aβ4-40, Aβ3pE-40, Aβ1-40. Correlations matrices generated for cored plaques showed a negative correlation for Aβx-40 (Aβ 1-40 Aβ3pE-40, Aβ3-40, Aβ4-40) with Aβx-42 species. In contrast different x-42 truncations showed positive correlations with each other, while x-40 peptides display positive correlation with other x-40 species (SI Fig. S3, Fig. 5E).

Most interestingly, the multivariate HCA image analysis **(Fig. 1)** identified a characteristic subtype of fibrillar plaques that show morphological distinct phenotypes reminiscent of what has previously described as coarse grain plaques (Boon *et al*. 2020). Investigation of plaque load for coarse grain (CG) plaques showed a significantly higher load of those plaques in fAD brain as compared to sAD (p=0.029) **(Fig. S5)**. The next aim was hence to investigate differences in Aβ signatures in between coarse grain and cored plaques both in fAD and sAD brain. For this OPLS DA models were generated for comparing the two plaque types within fAD and sAD. When comparing cored- and coarse grain plaques, OPLS DA showed a clear separation of in both fAD and in sAD **(Fig. 5A I,B I, SI Fig. S5a, b)**. A prominent but not similar strong separation was observed in between coarse grained plaques in fAD and sAD **(Fig S5c)**. The OPLS DA derived VIP, HCA heatmaps and univariate statistical analysis revealed that coarse grain plaques across both sAD and fAD show higher levels of Aβx-40 species including Aβ4-40, Aβ3pE-40, and Aβ1-40. In contrast, both in sAD and fAD, Aβx-42 species were higher in cored plaques as compared to coarse grain plaques Aβ11pE-42, Aβ5-42(p<0.05), Aβ3pE-40(p<0,0001), Aβ3pE-42(p<0.05), and Aβ1-42 (p<0.05).

When comparing only coarse grain plaques, the results showed relatively higher levels of pyroglutamated Aβ x-40 species (Aβ3pE-40, Aβ11pE-40) in sAD CG as compared to fAD CG **(Fig. 5d)**.

Regression analyses of peptide species in coarse grain plaques showed a significant, negative correlation of Aβ1-40, Aβ3pE-40, Aβ3-40 with Aβ(x-42). Several more N-terminal Aβx-40 truncations like Aβ4-40, Aβ5-40, Aβ8-40 showed a weaker negative correlation with Aβ(x-42) (Fig. 5E, SI Fig. S6 and S7)

### 6. Coarse grain plaques display amyloid signatures designative to CAA

The MS signatures observed for coarse grain plaques were dominated by Aβ1-40 species. We and others have previously described that vascular deposits (CAA) show Aβ1-40 dominated Aβ profiles as well. We therefore compared MALDI MSI Aβ patterns between CAA and CG plaques using OPLS-DA **(Fig.6A, B)**. The results show lower levels of Aβ1-40 and Aβ3pE-40 in CG plaques as compared to CAA. Conversely, Aβx-42 species including Aβ1-42, Aβ5-42, Aβ3pE-42, and Aβ11pE-42 were increased in CG plaques as compared to CAA **(Fig. 6C, D)**.

**Figure 6.**
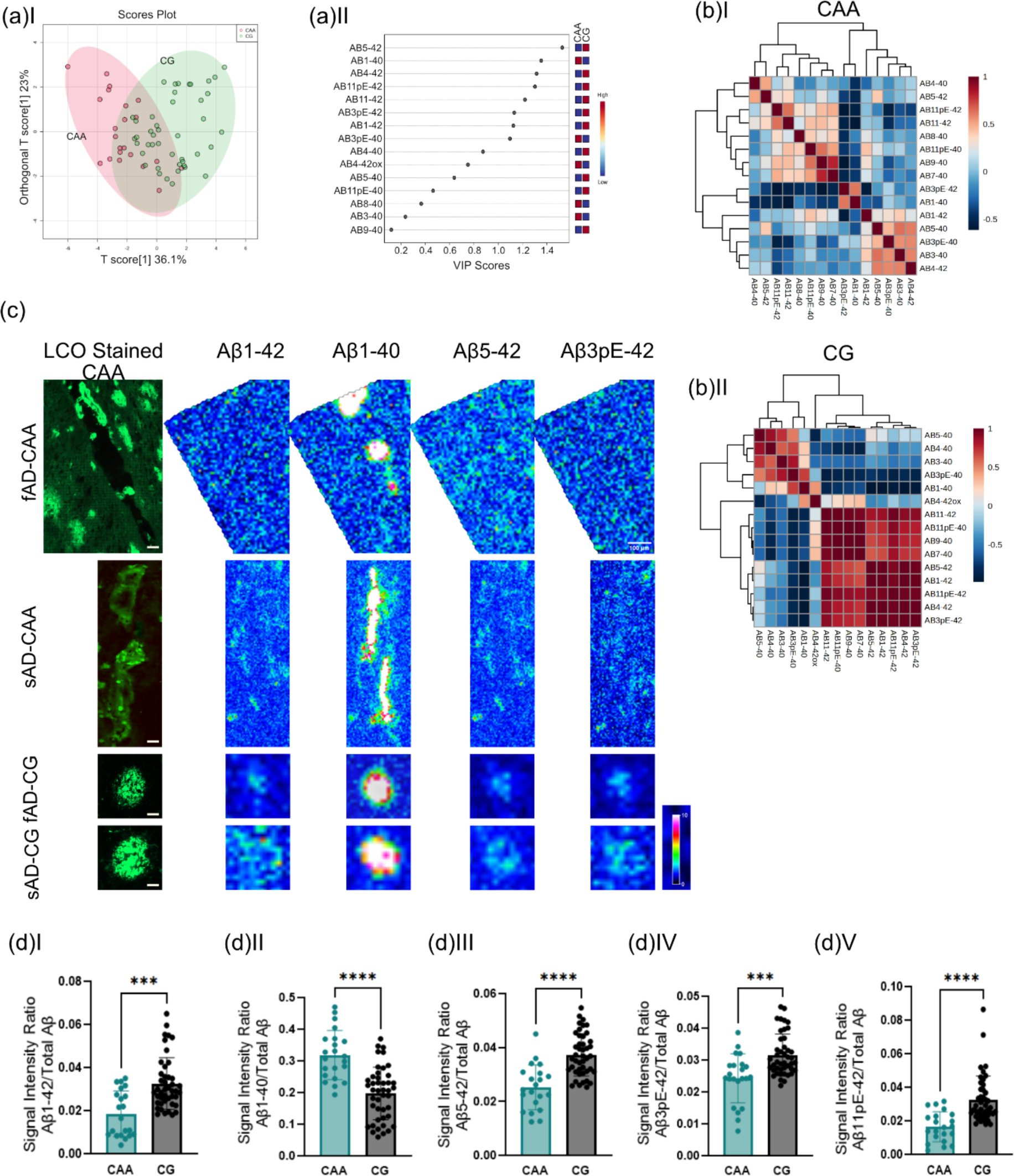
Coarse grain plaques display amyloid signatures designative to CAA. (a) OPLS-DA of CAA and CG plaques show partial separation (OPLS model characteristics: R2X-0.329; R2Y-0.439; Q2-0.402) and (b) VIP scores. (c) Single ion maps of different Aβ species. Scalebar = 50µm. (D) Univariate comparison between CAA and CG. *p<0.05, **p<0.01, ***p<0.001, ****p<0.0001 Number of patients n=8 (sAD), n=5 (fAD); number of plaques N=21 (CAA), N=43 (CG)

## Discussion

Aβ plaques exist in different morphotypes (Saido *et al*. 1995; Iwatsubo *et al*. 1994) that differ in terms of their fibrillar nature, chemical composition and how they interact with the surroundings. Factors contributing to the formation of different Aβ morphotypes are not very well understood yet highlighting the importance to increase our chemical understanding of these plaques and their associated diversity. Conventional immunostaining can help identify various plaque types, but the chemical composition of these plaques remains unknown. Herein, we delineated plaque polymorphism using a multimodal chemical imaging strategy that combines functional amyloid microscopy (Nyström *et al*. 2017) and MALDI mass spectrometry imaging. This strategy allowed us to identify distinct structural plaque morphotypes and to characterize the Aβ phenotype of these plaques beyond what is possible with established staining techniques. Specifically, we delineated various forms of diffuse- and fibrillar plaques as well as vascular deposits (CAA) across sporadic AD (sAD) and familial AD (fAD).

### [Diffuse plaques and core plaques]

We show that fibrillar plaques are characterized by prominent deposition of Aβx-40 peptide along with higher AβxpE-42 as compared to diffuse plaques. We and others have shown that cored plaques are characterized by Aβx-40 deposition (Michno *et al*. 2019b). However, as Aβx-40 peptides are less prone to aggregation, the prominence of AβxpE-42 in AD plaques could explain this discrepancy in that this represents a hydrophobic functionalization of Aβ1-42 (Michno *et al*. 2019b; Selkoe & Hardy 2016). Indeed, N-terminal functionalization of Aβ through enzymatic conversion to pyroglutamate alters the biophysical properties like increased hydrophobicity of the peptides. A pyroglutamate-modified Aβ3-x, i.e. Aβ3pE-x has been reported to be present in significant fractions in AD and exhibits accelerated aggregation kinetics, higher toxicity (Bayer 2022), increased β sheet stability (Nath *et al*. 2023) and resistance to degradation in in-vivo than a non-modified Aβ peptide (Dammers *et al*. 2015).

### [Neuritic plaques and Non-Neuritic plaques]

To further characterize cored deposits as outlined by LCO microscopy, we combined of IHC (using PHF-1) and MALDI MSI, which allowed to distinguish those plaques into neuritic- and non neuritic cored plaques in sAD, while only PHF-1-positive, i.e., neuritic, cored plaques were observed in fAD. MALDI MSI revealed higher levels of oxidized forms of Aβ1-42 (Aβ1-42ox) and Aβ2-42 in neuritic plaques compared to non-neuritic plaques. Conversely, there was no noticeable difference in the levels of Aβ1-40 or AβxpE-42 between the two plaque types. Previous research has suggested that CSF levels of N-terminal truncated peptides like Aβ2-42 could be a more reliable indicator of AD pathology than Aβ 1-40/1-42 levels in CSF {Bibl, 2012 #12}. Aβ2-42 has further been shown to accelerate amyloid aggregation and to be associated with neurodegenerative changes linked to Aβ plaques, indicating its potential role in triggering Aβ aggregation (Bibl *et al*. 2012).

### [Cored plaques and CG plaques]

Using chemical imaging with mass spectrometry further allowed us to identify a distinct fibrillar plaque population that has recently been described as coarse grain (CG) plaque (Boon *et al*. 2020). Consequently, these plaques have also been reported in the past as fibrillar plaques, primitive plaques, or burnt-out plaques. CG plaques stand out as a distinct population of plaques with their unique physical and chemical traits that exhibit dense fibrillar deposition (Boon *et al*. 2020). Typically, CG plaques are large circular plaques (diameter of ∼80µm) in comparison with typical cored plaques (diameter of ∼50µm). CG plaques are devoid of corona while cored plaques are often seen with circular corona around the dense central deposition, referred to as core of the plaque.

In our analysis, we found that CG plaques are predominantly composed of Aβ1-40 and Aβx-40 species, along with smaller proportions of Aβx-42. On the contrary, classic cored plaques exhibit relatively smaller amounts of Aβ1-40 deposition at the core but display prominent Aβx-42 patterns, i.e. a higher proportion of Aβ3pE-42 and Aβ11pE-42. Previous MALDI and IHC experiments on cored plaques have shown that in cored plaques, the core is predominantly composed of Aβ1-40, while the diffuse corona is composed of Aβx-42 fibrils (Boon *et al*. 2020{Michno, 2019 #7)}. The larger diameter of the CG plaques can be attributed to the dense deposition of Aβx-40, as Aβ1-40, a major constituent of CG plaques has an inherent ability to form thick large fibrils. On the contrary, although the core of the cored plaque is mainly composed of Aβ1-40, overall composition depicts higher proportions of Aβx-42 species, wherein Aβ1-42 is known to form thin fibrils as seen in the corona or in diffused plaques. Indeed, Aβ1-40 is a soluble peptide often observed in large proportions in CSF, which in pathological conditions are prone to aggregate and accumulate in dense, pre-seeded Aβ deposits. While Aβ1-42 is known for its sticky nature and its ability to seed and aggregate on its own. CG plaques showed further a high degree of Aβx-40 pyroglutamation. Similarly, to AβxpE-42, pyroglutamation of Aβx-40 species results in enhanced seeding ability, accelerated aggregation kinetics and increased neurotoxic effects (Bayer 2022). Pyroglutamate modification of Aβ1-40, together with N terminal truncation of Aβ1-40 into Aβ4-40, could be the contributing factors towards the formation of coarse grain plaques despite lower amounts of Aβ1-42 (Dunys *et al*. 2018; Schlenzig *et al*. 2009).

### [Neurotoxicity, Axonal and Dystrophic Neurites in Fibrillar Plaque Morphotypes]

In our analysis, CG plaques were observed in both sporadic and familial AD. Here, CG load was higher in fAD than in sAD. In line with previous data, CG plaques are more prominent in EOAD rather than in LOAD patients (Boon *et al*. 2020). Coarse grain plaques are observed in both APOE-e4 and -non e4 AD cases but are more prominently seen in homozygous e4 carriers (Boon *et al*. 2020). CG plaques further show increased levels of axonal and neuritic dystrophy, which is relevant as the load of neuritic, fibrillar plaques correlated with the clinical presentation of dementia. Both suggest that these plaques are the key neurotoxic amyloid entities driving disease pathogenesis.

Indeed, prolonged exposure of neurons to Aβ plaques induces neurotoxic changes often triggering stress-mediated response (REF). Such intra-neuronal changes could trigger, for example, an autophagy-related cascade of events and hyperphosphorylation of tau either at the synaptic end or along the axonal length of the neuron, leading to both synaptic and axonal dystrophy, which is visualized with Reticulon-3 (RTN3) (Sharoar *et al*. 2019) as well as paired helical filament 1 (PHF-1) (Otvos *et al*. 1994) (Su *et al*. 1996; Dickson 1999)

Fibrillar plaques like CP and CG observed in both sAD and fAD cases show significant neuritic pathology as compared to diffuse plaques (sAD) and cotton wool plaques (fAD), which further suggests that fibrillar plaques are advanced disease stage/neurotoxicity related plaque phenotypes. Indeed, the presence of Aβ plaques associated with dystrophic neurites is an indicator of late-stage AD, as tau-related changes in the neurites occur in the advanced stages of the disease (Dickson 1999; Dickson 2001), providing further support that CG plaques advanced-stage plaques.

Further, both cored and coarse grain plaques showed significantly higher levels of pyroglutamation of the respective dominating Aβ species (Aβx-42 and Aβx-40). This ties well together with results from clinical trials using Donanemab that targets N-terminal pyroglutamate Aβ present in those plaques (Schelle *et al*. 2017; Maia *et al*. 2013), further highlighting the neurotoxic relevance of those plaque phenotypes. Further, a more pronounced neuroinflammatory and reactive glial cell response was described for CG as compared to other plaque types. Together this indicates that CG formation may be associated with changes in neuroinflammatory response and neurovascular clearance. All these observations support the clinical relevance of CG plaque pathology in cerebral amyloidosis in AD. All these patterns suggest that CG plaques are a more mature plaque stage, more implicated with advanced stages of AD.

### [CG vs CAA]

Finally, our results show that the Aβ pattern observed for coarse grain plaques was generally very similar to that of CAA deposits. This is interesting as CAA load has also been associated with clinical dementia both in AD but also familial British and Danish dementia (Michno *et al*. 2022). Indeed, a vascular component in CG formation has previously been proposed, where CGs were found to display a characteristic increase in vascular and capillary pathology markers including norrin, laminin and collagen IV, respectively (Boon *et al*. 2020). In our findings, we observed increased levels of Aβx-42 in CG compared to CAA, indicating the significance of Aβx-42 in parenchymal plaque formation as opposed to passive Aβ1-40 deposition in the vasculature. Concurrently, higher levels of Aβ1-40 were noted in CAA compared to CG, implying a potential association with vascular permeability or leakage. Further, a more pronounced neuroinflammatory and reactive glial cell response was described for CG as compared to other plaque types. Together this indicates that CG formation may be associated with changes in neuroinflammatory response and neurovascular clearance. All these observations support the clinical relevance of CG plaque pathology in cerebral amyloidosis in AD. All these patterns suggest that CG plaques are a more mature plaque stage, more implicated with advanced stages of AD.

In summary, we demonstrate a novel chemical imaging paradigm to interrogate the biochemical signature of morphologically distinct amyloid plaque pathology in AD. Amyloid maturation has previously been implicated in AD pathogenesis proposing that the formation of fibrillar plaques is critical in AD pathology. The present data suggest that fibrillar plaques and more specifically coarse grain plaques represent a pathological signature indicative of both advanced AD. The results and methods described here are very relevant to AD research and AD drug development, particularly in the light of recent FDA approvals for novel biomedicines focusing on the reduction of plaque load in AD patients and supposedly target differentially modified and aggregated forms of Aβ. Given that the effect of those drugs has recently only been studied with established biochemical techniques, it is crucial to further understand what structured are targeted and inhibited by those drugs to refine those strategies and minimize adverse effects.

## Author contribution

JH conceived and designed the study. TL selected the cases from QSBB archives. SK, JG, DJ performed experiments. SK, JG, DJ, MD, WM, KB, TL, NR, HZ and JH analyzed and discussed the data. JH and SK wrote the manuscript.

## ACKNOWLEDGEMENTS

We thank the staff at Centre for Cellular Imaging (CCI), Core Facilities, The Sahlgrenska Academy, University of Gothenburg, for help with development of the hyperspectral imaging paradigm and microscopy expertise. The Queen Square Brain Bank is supported by the Reta Lila Weston Institute of Neurological Studies, UCL Queen Square Institute of Neurology.

## Abbreviations

AD: Alzheimer’s disease
Aβ: beta-amyloid
LCO: Luminescent Conjugated Oligothiophenes
MSI: Mass Spectrometry Imaging
ROI: region of Interest

## Supporting Information

**SI Table S1:** Masses of the detected Aβ isoforms.

| Peptide | Peptide Sequence | [M+H]<sup>+</sup> | [M+H]<sup>+</sup> Avg |
| --- | --- | --- | --- |
| Aβ11pE-40 | pEVHHQKLVFFAEDVGSNKGAIIGLMVGGVV | 3132.67684 | 3134 |
| Aβ11pE-42 | pEVHHQKLVFFAEDVGSNKGAIIGLMVGGVVIA | 3317.79802 | 3319 |
| Aβ11-42 | EVHHQKLVFFAEDVGSNKGAIIGLMVGGVVIA | 3334.79802 | 3336 |
| Aβ9-40 | GYEVHHQKLVFFAEDVGSNKGAIIGLMVGGVV | 3370.76164 | 3372 |
| Aβ8-40 | SGYEVHHQKLVFFAEDVGSNKGAIIGLMVGGVV | 3457.79366 | 3459 |
| Aβ7-40 | DSGYEVHHQKLVFFAEDVGSNKGAIIGLMVGGVV | 3572.82061 | 3574 |
| Aβ4-40 | FRHDSGYEVHHQKLVFFAEDVGSNKGAIIGLMVGGVV | 4013.04904 | 4012.042 |
| Aβ5-42 | RHDSGYEVHHQKLVFFAEDVGSNKGAIIGLMVGGVVIA | 4050.10181 | 4052 |
| Aβ3pE-40 | pEFRHDSGYEVHHQKLVFFAEDVGSNKGAIIGLMVGGVV | 4124.09164 | 4127 |
| Aβ4-42 | FRHDSGYEVHHQKLVFFAEDVGSNKGAIIGLMVGGVVIA | 4197.17022 | 4199.5 |
| Aβ3pE-42 | pEFRHDSGYEVHHQKLVFFAEDVGSNKGAIIGLMVGGVVIA | 4308.21281 | 4311 |
| Aβ1-40 | DAEFRHDSGYEVHHQKLVFFAEDVGSNKGAIIGLMVGGVV | 4328.15569 | 4329.5 |
| Aβ2-42 | AEFRHDSGYEVHHQKLVFFAEDVGSNKGAIIGLMVGGVVIA | 4397.24993 | 4400 |
| Aβ1-42 | DAEFRHDSGYEVHHQKLVFFAEDVGSNKGAIIGLMVGGVVIA | 4512.27687 | 4515.5 |
| Aβ1-42ox | DAEFRHDSGYEVHHQKLVFFAEDVGSNKGAIIGLMVGGVVIA +oxidation | 4528.27687 | 4531.5 |

**Figure S1.**
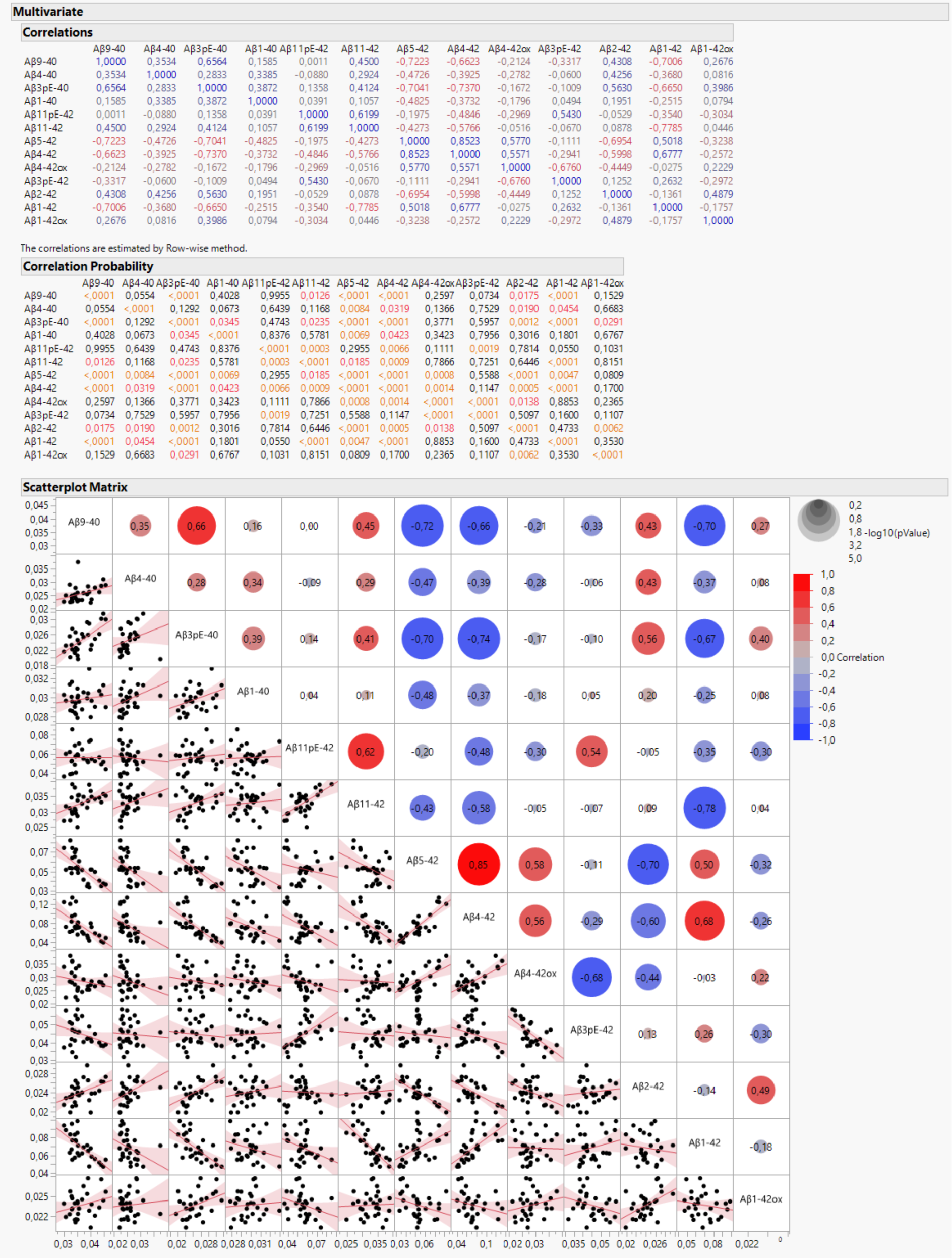
Correlation analysis of amyloid peptides in sAD-Core Plaque

**Figure S2.**
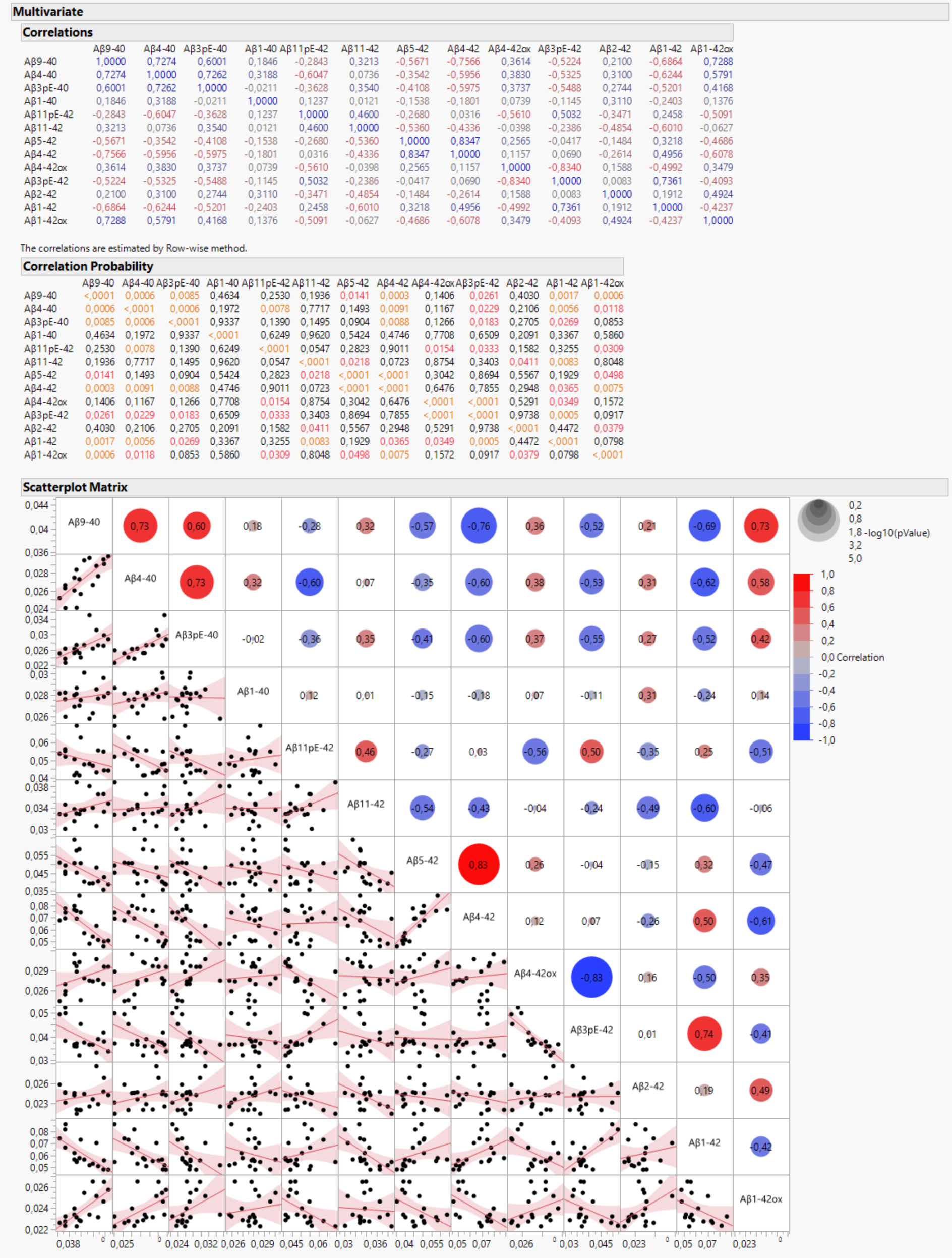
Correlation analysis of amyloid peptides in sAD-Diffused Plaque

**Figure S3.**
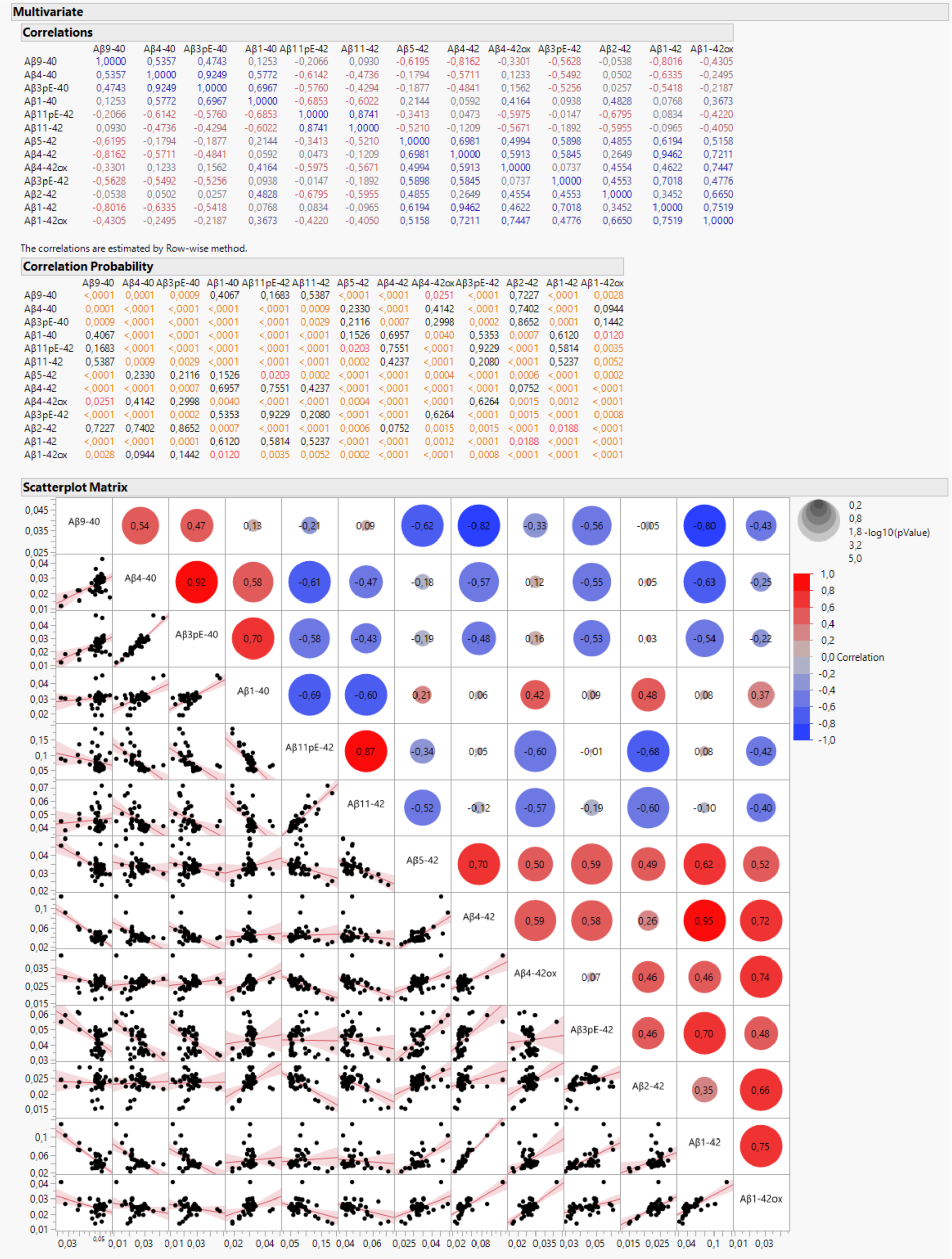
Correlation analysis of Aβ peptides in fAD-Core Plaque

**Figure S4.**
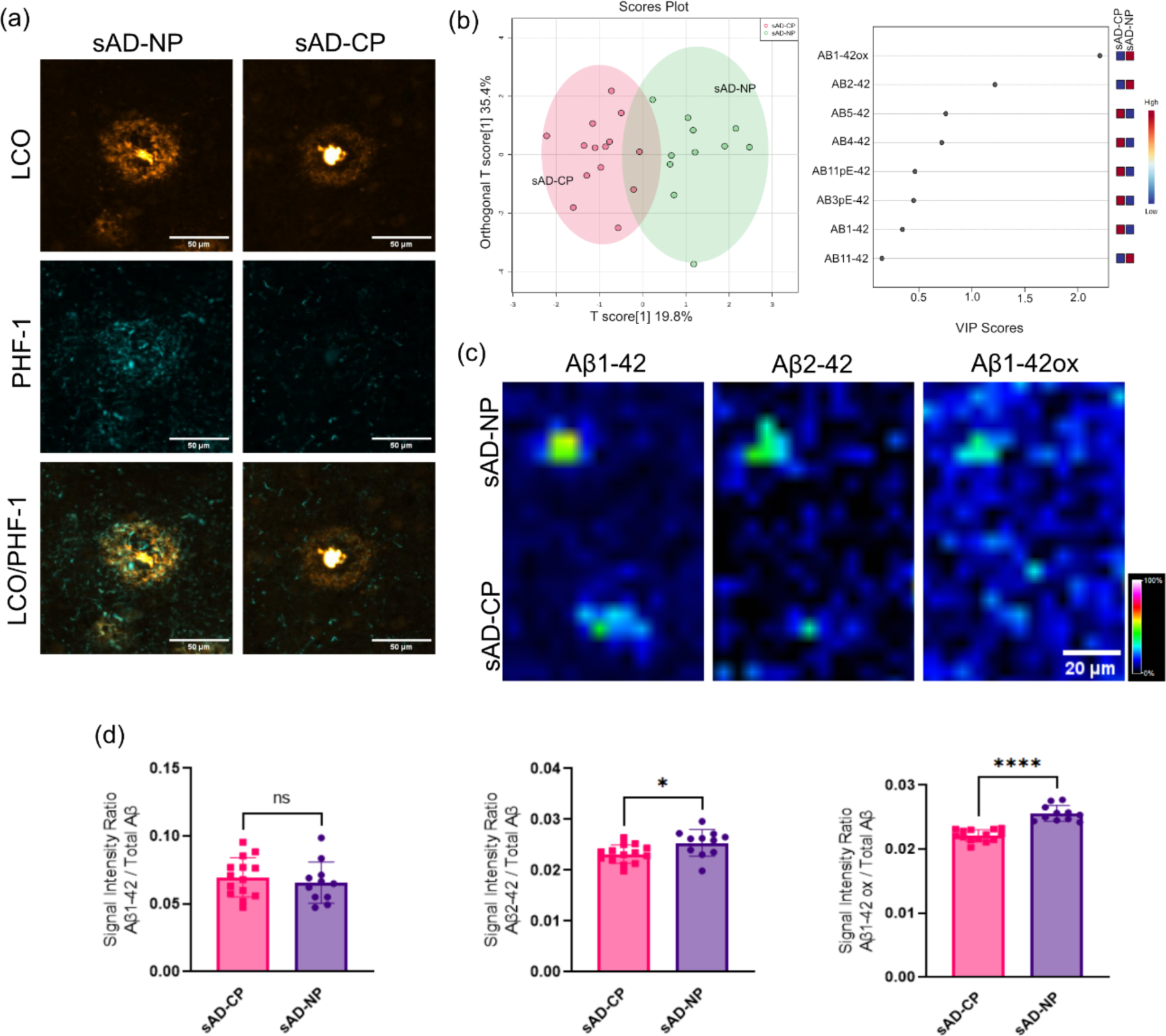
MALDI signatures of neuritic plaques. (a) Delineating plaque types by means of fluorescent microscopy using LCO amyloid probes (q and h FTAA) along with PHF-1 IHC. Neuritic plaques show higher levels of PHF-1 Tau positive neurites. (b) OPLS DA of MALDI signatures(OPLS model characteristics: R2X-0.256; R2Y-0.558; Q2-0.486). Here the VIP reveal elevated levels of Aβ2-42 and Aβ1-42ox in neuritic plaques as compared to cored plaques. (c, d) Bar graphs and single ion images. Scale bar: 30um. Intensity scale: rel. intensity in %.

**Figure S5.**
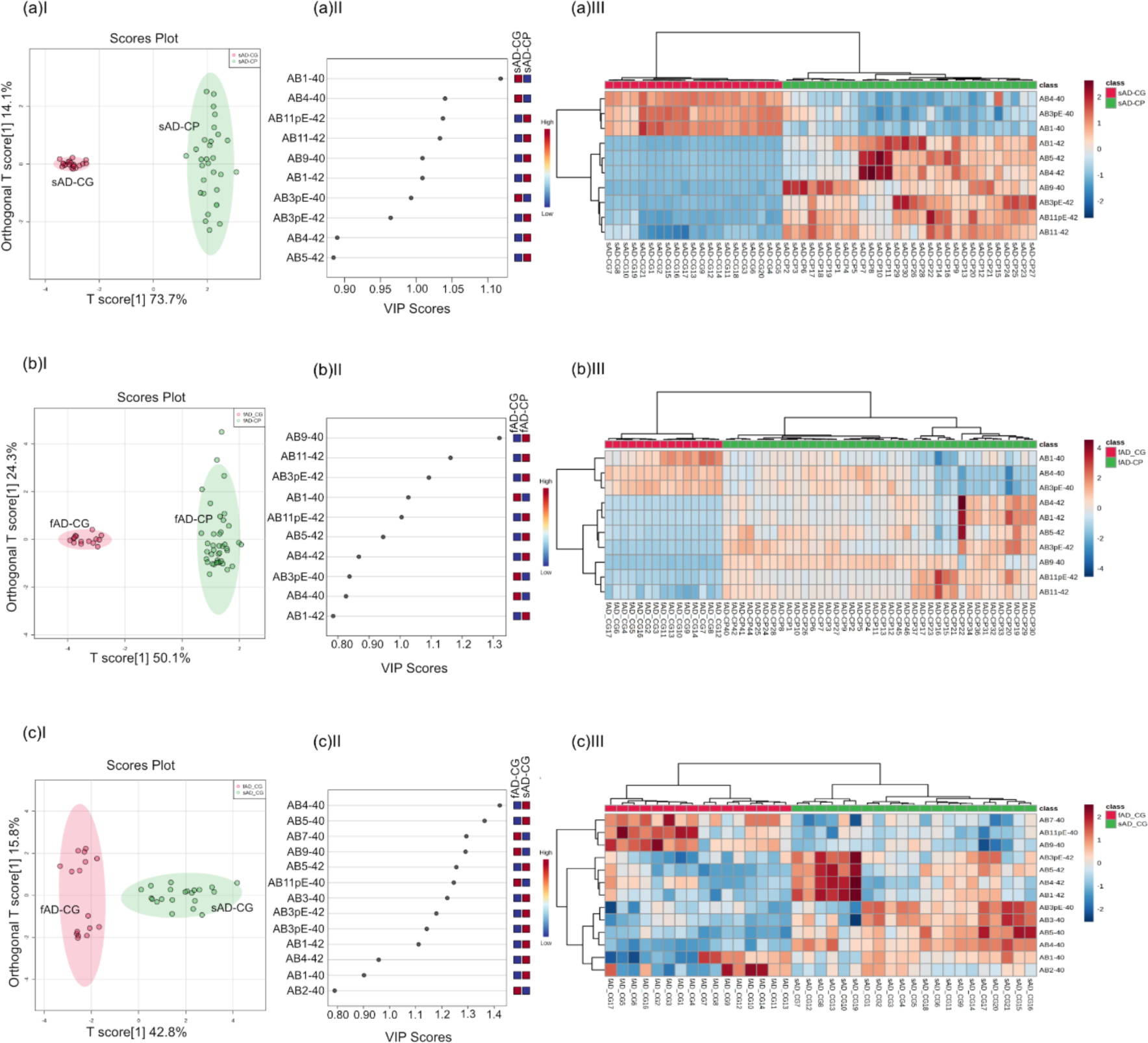
Comparative analysis of Aβ patterns in cored and coarse grain plaques. (a) sAD cored plaques vs coarse grain plaques (OPLS model characteristics: R2X-0.737; R2Y-0.948; Q2-0.947)(b) fAD cored plaques vs coarse grain plaques (OPLS model characteristics: R2X-0.501; R2Y-0.813; Q2-0.81) (c) sAD coarse grain plaques vs fAD coarse grain plaques (OPLS model characteristics: R2X-0.428; R2Y-0.852; Q2-0.835) (a-cI) OPLS-DA score plot. (a-cII) VIP scores. (a-cIII) HCA heatmap.

**Figure S6.**
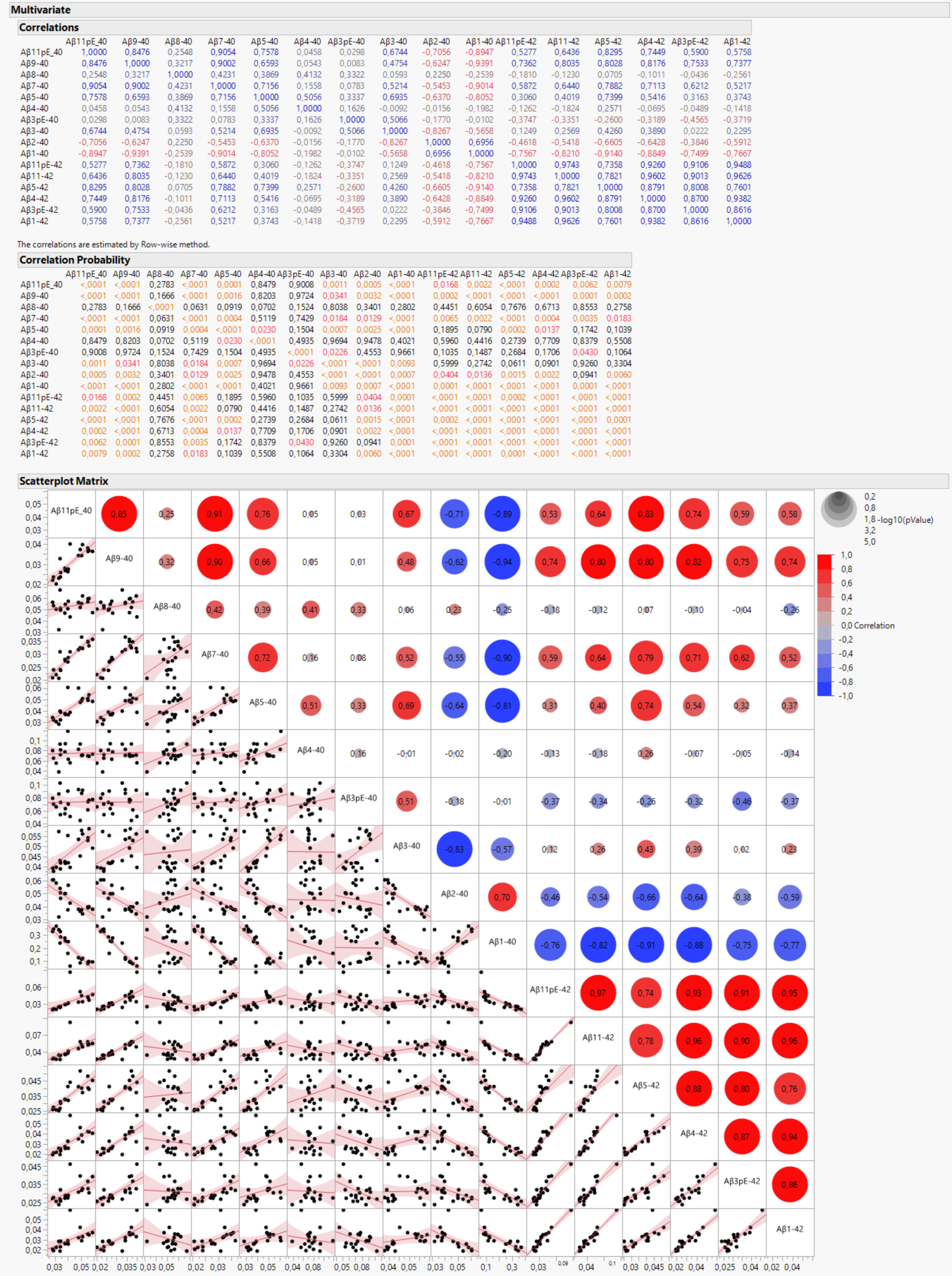
Correlation analysis of amyloid peptides in fAD-Coarse Grain

**Figure S7.**
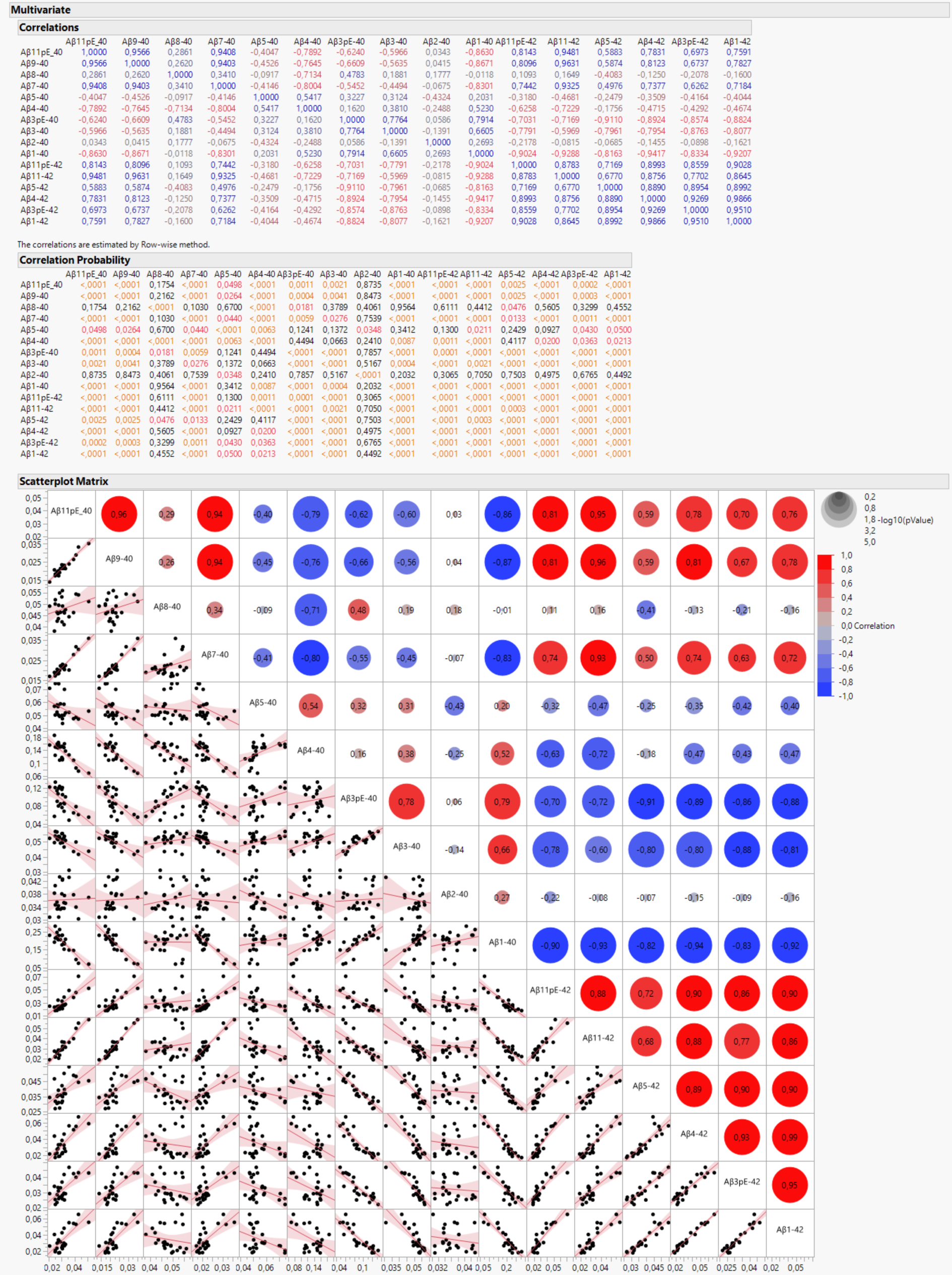
Correlation analysis of amyloid peptides in sAD-Coarse Grain

## References

1. Bayer, T. A. (2022) Pyroglutamate Aβ cascade as drug target in Alzheimer’s disease. Mol Psychiatry 27, 1880–1885.

2. Bibl, M., Gallus, M., Welge, V., Esselmann, H., Wolf, S., Rüther, E. and Wiltfang, J. (2012) Cerebrospinal fluid amyloid-β 2-42 is decreased in Alzheimer’s, but not in frontotemporal dementia. J Neural Transm (Vienna) 119, 805–813.

3. Boon, B. D. C., Bulk, M., Jonker, A. J. et al. (2020) The coarse-grained plaque: a divergent Aβ plaque-type in early-onset Alzheimer’s disease. Acta neuropathologica 140, 811–830.

4. Braak, H. and Braak, E. (1991) Neuropathological stageing of Alzheimer-related changes. Acta neuropathologica 82, 239–259.

5. Carlred, L., Michno, W., Kaya, I., Sjovall, P., Syvanen, S. and Hanrieder, J. (2016) Probing amyloid-beta pathology in transgenic Alzheimer’s disease (tgArcSwe) mice using MALDI imaging mass spectrometry. J Neurochem 138, 469–478.

6. Dammers, C., Gremer, L., Reiß, K. et al. (2015) Structural Analysis and Aggregation Propensity of Pyroglutamate Aβ(3-40) in Aqueous Trifluoroethanol. PLoS One 10, e0143647.

7. Dickson, D. W. (1999) Neuropathologic differentiation of progressive supranuclear palsy and corticobasal degeneration. Journal of Neurology 246, II6-II15.

8. Dickson, D. W. (2001) Neuropathology of Alzheimer’s disease and other dementias. Clinics in Geriatric Medicine 17, 209–228.

9. Dickson, T. C. and Vickers, J. C. (2001) The morphological phenotype of beta-amyloid plaques and associated neuritic changes in Alzheimer’s disease. Neuroscience 105, 99–107.

10. Dunys, J., Valverde, A. and Checler, F. (2018) Are N- and C-terminally truncated Aβ species key pathological triggers in Alzheimer’s disease? J Biol Chem 293, 15419–15428.

11. EISAI (2022) Lecanemab confirmatory Phase 3 Clarity AD study met primary endpoint, showing highly statistically significant reduction of clinical decline in large global clinical study of 1,795 participants with early Alzheiemr’s disease. Eisai Inc., https://eisai.mediaroom.com/2022-09-27-LECANEMAB-CONFIRMATORY-PHASE-3-CLARITY-AD-STUDY-MET-PRIMARY-ENDPOINT,-SHOWING-HIGHLY-STATISTICALLY-SIGNIFICANT-REDUCTION-OF-CLINICAL-DECLINE-IN-LARGE-GLOBAL-CLINICAL-STUDY-OF-1,795-PARTICIPANTS-WITH-EARLY-ALZHEIMERS-DISEASE.

12. Hanrieder, J., Ljungdahl, A., Falth, M., Mammo, S. E., Bergquist, J. and Andersson, M. (2011) L-DOPA-induced dyskinesia is associated with regional increase of striatal dynorphin peptides as elucidated by imaging mass spectrometry. Molecular & cellular proteomics : MCP 10, M111.009308.

13. Hardy, J. (1992) An anatomical cascade hypothesis for Alzheimer’s disease. Trends in Neurosciences 15, 200–201.

14. Hardy, J. and Selkoe, D. J. (2002) Medicine - The amyloid hypothesis of Alzheimer’s disease: Progress and problems on the road to therapeutics. Science 297, 353–356.

15. Hayato, S., Takenaka, O., Sreerama Reddy, S. H., Landry, I., Reyderman, L., Koyama, A., Swanson, C., Yasuda, S. and Hussein, Z. (2022) Population pharmacokinetic-pharmacodynamic analyses of amyloid positron emission tomography and plasma biomarkers for lecanemab in subjects with early Alzheimer’s disease. CPT Pharmacometrics Syst Pharmacol.

16. Iwatsubo, T., Odaka, A., Suzuki, N., Mizusawa, H., Nukina, N. and Ihara, Y. (1994) Visualization of A beta 42(43) and A beta 40 in senile plaques with end-specific A beta monoclonals: evidence that an initially deposited species is A beta 42(43). Neuron 13, 45–53.

17. Kirmess, K. M., Meyer, M. R., Holubasch, M. S. et al. (2021) The PrecivityAD test: Accurate and reliable LC-MS/MS assays for quantifying plasma amyloid beta 40 and 42 and apolipoprotein E proteotype for the assessment of brain amyloidosis. Clin Chim Acta 519, 267–275.

18. Maia, L. F., Kaeser, S. A., Reichwald, J., Hruscha, M., Martus, P., Staufenbiel, M. and Jucker, M. (2013) Changes in amyloid-β and Tau in the cerebrospinal fluid of transgenic mice overexpressing amyloid precursor protein. Sci Transl Med 5, 194re192.

19. Matthews, K. A., Xu, W., Gaglioti, A. H., Holt, J. B., Croft, J. B., Mack, D. and McGuire, L. C. (2019) Racial and ethnic estimates of Alzheimer’s disease and related dementias in the United States (2015-2060) in adults aged ≥65 years. Alzheimers Dement 15, 17–24.

20. Michno, W., Koutarapu, S., Camacho, R. et al. (2022) Chemical traits of cerebral amyloid angiopathy in familial British-, Danish-, and non-Alzheimer’s dementias. J Neurochem 163, 233–246.

21. Michno, W., Nystrom, S., Wehrli, P. et al. (2019a) Pyroglutamation of amyloid-betax-42 (Abetax-42) followed by Abeta1-40 deposition underlies plaque polymorphism in progressing Alzheimer’s disease pathology. The Journal of biological chemistry.

22. Michno, W., Nyström, S., Wehrli, P. et al. (2019b) Pyroglutamation of amyloid-βx-42 (Aβx-42) followed by Aβ1-40 deposition underlies plaque polymorphism in progressing Alzheimer’s disease pathology. J Biol Chem 294, 6719–6732.

23. Michno, W., Stringer, K. M., Enzlein, T. et al. (2021) Following spatial Aβ aggregation dynamics in evolving Alzheimer’s disease pathology by imaging stable isotope labeling kinetics. Science Advances 7, eabg4855.

24. Michno, W., Wehrli, P., Meier, S. R., Sehlin, D., Syvänen, S., Zetterberg, H., Blennow, K. and Hanrieder, J. (2020) Chemical imaging of evolving amyloid plaque pathology and associated Aβ peptide aggregation in a transgenic mouse model of Alzheimer’s disease. J Neurochem 152, 602–616.

25. Michno, W., Wehrli, P. M., Blennow, K., Zetterberg, H. and Hanrieder, J. (2019c) Molecular imaging mass spectrometry for probing protein dynamics in neurodegenerative disease pathology. J Neurochem 151, 488–506.

26. Mintun, M. A., Lo, A. C., Duggan Evans, C. et al. (2021) Donanemab in Early Alzheimer’s Disease. N Engl J Med 384, 1691–1704.

27. Mirra, S. S., Heyman, A., McKeel, D. et al. (1991) The Consortium to Establish a Registry for Alzheimer’s Disease (CERAD). Part II. Standardization of the neuropathologic assessment of Alzheimer’s disease. Neurology 41, 479–486.

28. Montine, T. J., Phelps, C. H., Beach, T. G. et al. (2012) National Institute on Aging-Alzheimer’s Association guidelines for the neuropathologic assessment of Alzheimer’s disease: a practical approach. Acta neuropathologica 123, 1–11.

29. Nath, S., Buell, A. K. and Barz, B. (2023) Pyroglutamate-modified amyloid β(3-42) monomer has more β-sheet content than the amyloid β(1-42) monomer. Phys Chem Chem Phys 25, 16483–16491.

30. Nyström, S., Bäck, M., Nilsson, K. P. R. and Hammarström, P. (2017) Imaging Amyloid Tissues Stained with Luminescent Conjugated Oligothiophenes by Hyperspectral Confocal Microscopy and Fluorescence Lifetime Imaging. J Vis Exp.

31. Nystrom, S., Psonka-Antonczyk, K. M., Ellingsen, P. G. et al. (2013) Evidence for age-dependent in vivo conformational rearrangement within Abeta amyloid deposits. ACS Chem Biol 8, 1128–1133.

32. Otvos, L., Jr., Feiner, L., Lang, E., Szendrei, G. I., Goedert, M. and Lee, V. M. (1994) Monoclonal antibody PHF-1 recognizes tau protein phosphorylated at serine residues 396 and 404. J Neurosci Res 39, 669–673.

33. Rafii, M. S., Sperling, R. A., Donohue, M. C. et al. (2022) The AHEAD 3-45 Study: Design of a prevention trial for Alzheimer’s disease. Alzheimers Dement.

34. Rasmussen, J., Mahler, J., Beschorner, N. et al. (2017) Amyloid polymorphisms constitute distinct clouds of conformational variants in different etiological subtypes of Alzheimer’s disease. Proc Natl Acad Sci U S A 114, 13018–13023.

35. Ryan, N. S., Nicholas, J. M., Weston, P. S. J. et al. (2016) Clinical phenotype and genetic associations in autosomal dominant familial Alzheimer’s disease: a case series. Lancet Neurol 15, 1326–1335.

36. Saido, T. C., Iwatsubo, T., Mann, D. M., Shimada, H., Ihara, Y. and Kawashima, S. (1995) Dominant and differential deposition of distinct beta-amyloid peptide species, A beta N3(pE), in senile plaques. Neuron 14, 457–466.

37. Satlin, A., Wang, J., Logovinsky, V., Berry, S., Swanson, C., Dhadda, S. and Berry, D. A. (2016) Design of a Bayesian adaptive phase 2 proof-of-concept trial for BAN2401, a putative disease-modifying monoclonal antibody for the treatment of Alzheimer’s disease. Alzheimers Dement (N Y) 2, 1–12.

38. Schelle, J., Häsler, L. M., Göpfert, J. C. et al. (2017) Prevention of tau increase in cerebrospinal fluid of APP transgenic mice suggests downstream effect of BACE1 inhibition. Alzheimers Dement 13, 701–709.

39. Schlenzig, D., Manhart, S., Cinar, Y., Kleinschmidt, M., Hause, G., Willbold, D., Funke, S. A., Schilling, S. and Demuth, H. U. (2009) Pyroglutamate formation influences solubility and amyloidogenicity of amyloid peptides. Biochemistry 48, 7072–7078.

40. Schmidt, M. L., Robinson, K. A., Lee, V. M. and Trojanowski, J. Q. (1995) Chemical and immunological heterogeneity of fibrillar amyloid in plaques of Alzheimer’s disease and Down’s syndrome brains revealed by confocal microscopy. Am J Pathol 147, 503–515.

41. Selkoe, D. J. (2002) Alzheimer’s disease is a synaptic failure. Science 298, 789–791.

42. Selkoe, D. J. and Hardy, J. (2016) The amyloid hypothesis of Alzheimer’s disease at 25 years. EMBO Mol Med 8, 595–608.

43. Sharoar, M. G., Hu, X., Ma, X. M., Zhu, X. and Yan, R. (2019) Sequential formation of different layers of dystrophic neurites in Alzheimer’s brains. Mol Psychiatry 24, 1369–1382.

44. Sims, J. R., Zimmer, J. A., Evans, C. D. et al. (2023) Donanemab in Early Symptomatic Alzheimer Disease: The TRAILBLAZER-ALZ 2 Randomized Clinical Trial. JAMA 330, 512–527.

45. Su, J. H., Cummings, B. J. and Cotman, C. W. (1996) Plaque biogenesis in brain aging and Alzheimer’s disease: I. Progressive changes in phosphorylation states of paired helical filaments and neurofilaments. Brain Research 739, 79–87.

46. Swanson, C. J., Zhang, Y., Dhadda, S. et al. (2021) A randomized, double-blind, phase 2b proof-of-concept clinical trial in early Alzheimer’s disease with lecanemab, an anti-Abeta protofibril antibody. Alzheimers Res Ther 13, 80.

47. Thal, D. R., Rüb, U., Orantes, M. and Braak, H. (2002) Phases of A beta-deposition in the human brain and its relevance for the development of AD. Neurology 58, 1791–1800.

48. Thal, D. R., Walter, J., Saido, T. C. and Fändrich, M. (2015) Neuropathology and biochemistry of Aβ and its aggregates in Alzheimer’s disease. Acta Neuropathol 129, 167–182.

49. Timmers, M., Tesseur, I., Bogert, J. et al. (2019) Relevance of the interplay between amyloid and tau for cognitive impairment in early Alzheimer’s disease. Neurobiol Aging 79, 131–141.

50. van Dyck, C. H., Swanson, C. J., Aisen, P. et al. (2023) Lecanemab in Early Alzheimer’s Disease. N Engl J Med 388, 9–21.

